# Whole-genome resequencing reveals the population structure, genomic diversity, and demographic history of American chestnut (*Castanea dentata*)

**DOI:** 10.1101/2022.02.11.480151

**Authors:** Alexander M Sandercock, Jared W Westbrook, Qian Zhang, Hayley A Johnson, Thomas M Saielli, John A Scrivani, Sara F Fitzsimmons, Kendra Collins, Jeremy Schmutz, Jane Grimwood, Jason A Holliday

## Abstract

American chestnut (*Castanea dentata*) was once the most economically and ecologically important hardwood species in the United States. In the first half of the 20th century, an exotic fungal pathogen – *Cryphonectria parasitica* – decimated the species, killing approximately four billion trees. Two approaches to developing blight resistant American chestnut populations show promise, but both will require introduction of adaptive genomic diversity from wild germplasm to produce diverse, locally adapted reforestation populations. Here we characterize population structure, demographic history, and genomic diversity in a range-wide sample of 384 wild American chestnuts to inform conservation and breeding with blight resistant varieties. Population structure analyses with DAPC and ADMIXTURE suggest that the chestnut range can be roughly divided into northeast, central, and southwest populations. Within-population genomic diversity estimates revealed a clinal pattern with the highest diversity in the southwest, which likely reflects bottleneck events associated with Quaternary glaciation. Finally, we identified genomic regions under positive selection within each population, which suggests that defense against fungal pathogens is a common target of selection across all populations. Taken together, these results show that American chestnut underwent a postglacial expansion from the southern portion of its range leading to three extant populations. These populations will serve as management units for breeding adaptive genetic variation into the blight-resistant tree populations for targeted reintroduction efforts.

## 1 Introduction

The American chestnut (*Castanea dentata* (Marsh.) Borkh) is a deciduous tree with a widespread historical range in the eastern United States and southeastern Canada (1, 2). Historical records describe *C. dentata* as a canopy hardwood tree that typically grew 60 – 90 feet high, but that could exceed 120 feet, with a diameter of 3-5 feet (3, 4). The rapid growth of American chestnut, coupled with its decay resistant wood, previously made it the single most valuable hardwood species in the United States (5, 6). Moreover, the prodigious and reliable seed crop was an important source of food and feed throughout its native range (7).

In the first half of the 20th century, an exotic fungal pathogen *— Cryphonectria parasitica* – decimated the American chestnut, killing billions trees. While >400 million American chestnuts survive in the forests of the eastern United States (8), the vast majority of these are collar sprouts from trees that germinated before arrival of the blight (9, 10). Although some of these sprouts are able to flower before being killed by blight, most are reinfected before they are able to produce nuts. The American chestnut is thus considered functionally extinct.

Successful restoration of the American chestnut will rely on an accurate understanding of extant genomic diversity in the wild (11), which can be leveraged to increase effective population size and adaptability of blight-resistant populations (12). The ultimate goal of this project is to prioritize geographic areas for *ex situ* conservation through propagation of wild trees. These trees will then be used to introgress natural genetic variation into blight resistant populations. The first step in this process is to define broad management units on the basis of population structure and postglacial history for the species (13).

Previous population genetic studies used microsatellite and ddRAD-seq to characterize the population structure and genetic diversity of American chestnut (14–17). These studies suggest the southern portions of the American chestnut range are the most genetically diverse and that two populations exist: either a distinct northeastern population (15, 16) or a distinct southern population (17). Furthermore, these studies suggest that the American chestnut underwent a postglacial migration northward from a southern glacial refugium (15, 17). While these studies provide our first glimpse of the extant patterns of genetic diversity in chestnut, they are limited by relatively low geographic sampling density and genotyping methods that may lack power to comprehensively characterize genome-wide diversity and signatures of selection (18, 19).

In this study, we used ≈ 17X whole-genome resequencing (WGS) data for each of 384 American chestnut genotypes, sampled from across the entire historical species range, to (i) estimate the population structure; (ii) describe the demographic history and contemporary barriers to gene flow; (iii) evaluate genomic diversity within each population; (iv) and identify signatures of selection within this iconic species. To our knowledge, our use of WGS is the first for American chestnut and allows the use of modern demographic inference techniques that were inaccessible to previous studies. Wholegenome resequencing captures many times more variant sites and has improved statistical accuracy than other low-density methods (20). Working in collaboration with The American Chestnut Foundation (TACF), the results of this study will be used to identify management units for germplasm conservation, and will aid in the overall efforts to restore the American chestnut population to its pre-blight abundance in the eastern US.

## 2 Materials and Methods

### 2.1. Leaf sample collection

TACF staff and volunteers sampled leaves from ≈1000 American chestnut trees throughout the species range. The youngest leaves were sampled from each tree in May through July over three growing seasons (2018 – 2020). During and immediately after collection, leaves were kept cool with ice and refrigeration or were desiccated and stored with silica gel. Leaves were shipped to Virginia Tech within 3 weeks of collection and stored at -80°C. Coordinates of the sampled trees were retrieved from TACF’s dentataBase (www.acf.herokuapp.com) and TreeSnap (21)(https://treesnap.org/).

### 2.2. DNA isolation and sequencing

From a cohort of ≈1,000 available leaf samples, we selected 384 genotypes for WGS (Fig. 1) on the basis of DNA quality and geography, with the goal of including as much of the historical range as possible. Leaves were ground to a fine powder using a Spex 2000 Geno/Grinder and ceramic beads with three 45 second intervals of grinding interspersed by submersion in liquid nitrogen. DNA was extracted with Qiagen’s DNAeasy Plant DNA extraction kit. For samples 1-96, a modified phenolchloroform cleanup step was performed to remove organic contaminants from older leaves. For samples 97 through 384, an additional 100% ethanol wash step was performed to remove organic contaminants that carried over from the previous steps and to dry the spin column membrane. DNA quality and concentration were measured by a Nanodrop One^c^ and Qubit 3.0 Fluorometer, respectively. When DNA concentration was low, a secondary CTAB-based extraction was used. DNA was stored in a 100-200 µl AE solution in a - 20°C freezer. PCR-free library preparation and genomic sequencing were conducted at the HudsonAlpha Institute for Biotechnology. Libraries were sequenced in batches of 48 on an Illumina NovaSeq 6000 instrument in 2×150bp paired-end mode with a target depth of 20x.

**Fig. 1.**
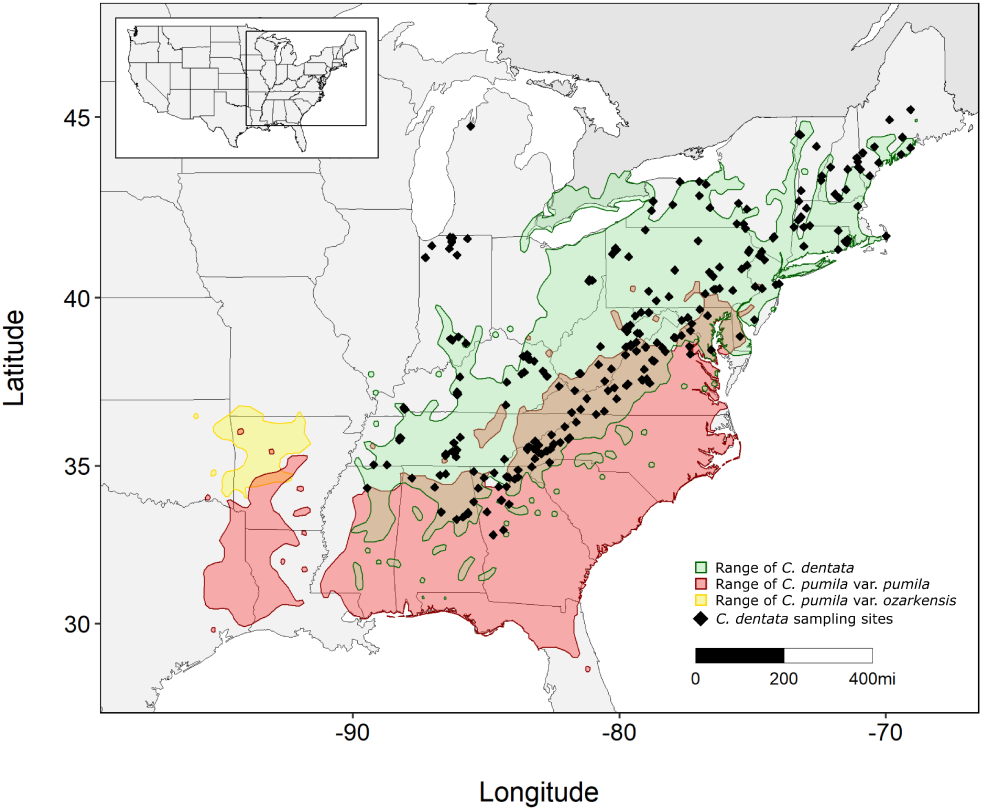
Range map of North American *Castanea* species and locations of the se- lected 384 American chestnut samples.

We also performed WGS on a *Castanea* species reference panel of 95 individuals to detect potential hybrid ancestry in the putative *C. dentata* samples. The reference panel included 19 *C. sativa*, 15 *C. pumila* var. *pumila*, 10 *C. pumila* var. *ozarkensis*, six *C. pumila* var. *alabamensis*, four *C. dentata*, one *C. seguinii*, two *C. henryi*, 18 *C. crenata*, and 20 *C. mollissima*. Leaf samples from Asian *Castanea* species (*C. mollissima, C. crenata*, and *C. henryi*) were collected from trees planted in the U.S. DNA from *C. sativa* was provided by R. Costa from trees in Portugal. *C. pumila* samples were collected from native locations (non-planted) in the U.S. DNA was extracted as above. Sequencing was performed at the Duke University Center for Genomic and Computational Biology, where libraries were prepared with an Illumina Nextera kit and sequenced in an Illumina NovaSeq 6000 S4 flowcell (48 samples per lane) in 2×150bp paired-end mode.

### 2.3. Bioinformatics

Bioinformatic analyses were performed on Virginia Tech’s Advanced Research Computing (ARC) system. SNPs were called using a custom pipeline adapted from the Broad Institute’s Genome Analysis Toolkit (GATK v3.8) best practices (22). Individual fastq files were aligned using the Burrows-Wheeler Aligner (BWA) mem algorithm with the -M and -R flags and the *C. dentata* genome as a reference (*Castanea dentata* v1.1; http://phytozome-next.jgi.doe.gov/). The resulting SAM files were converted to BAM format, sorted, and indexed using SAMtools (23, 24). If samples were sequenced on multiple lanes, the individual lanes for each sample were then combined into a single BAM file using samtools merge. For the *Castanea* species reference, where PCR was performed in the library preparation, PCR duplicates were removed from the BAM files using MarkDuplicates in picard.

The GATK HaplotypeCaller algorithm (25, 26) was used to call SNPs and INDELs by chromosome. Chromosome GVCF files for each individual were combined with GatherVcfs and individual GVCFs were merged with GenotypeGVCFs. Polymorphisms were quality filtered using the GATK VariantFiltration algorithm following GATK best practices. The VCF file was further filtered in VCFtools v0.1.17 (27) to include only biallelic SNPs (–max alleles 2) and SNPs with <10% missing data per site (–max-missing 0.9). Overall missing data per individual was checked in VCFtools (—missing-indv), and individuals with >10% missing data were removed. Unless otherwise noted, a minor allele frequency (MAF) filter was applied (MAF = 1/2n, where n = number of samples).

VCFtools was used to calculate the sequencing coverage and depth for the filtered VCF file. Genome-wide scans for SNP density were determined using the –SNPdensity option with 50kb windows. The mean sequencing depth per variant was calculated using the –site-mean-depth option and the mean sequencing depth per sample was determined using the – depth option. A smoothed line of the SNP density results for each chromosome was visualized in *ggplot2* using the default ‘gam’ parameter in geom_smooth.

### 2.4. Species identification and estimation of hybridization in wild American chestnut populations

We used the *Castanea* species reference dataset to test for evidence of introgression in our wild *C. dentata* samples. The *C. seguinii* and *C. henryi* samples were excluded as there were only one and two samples of these species, respectively. The VCF files from the *C. dentata* and species reference datasets were combined using bcftools merge and subsequently filtered with bcftools to retain biallelic SNPs and remove singleton SNPs (MAF = 1/2n, where n = number of samples) (28). Both SNPs and INDELs were retained for these analyses. ADMIXTURE was run with the –cv flag enabled to perform a five-fold cross-validation for K = 1-9 (29). The value of K was chosen as the most likely number of clusters when each *Castanea* species first separated into at least one distinct group (*C. pumila* varieties were considered as a single cluster). Putative *C. dentata* samples were classified as hybrids or misidentified if they had cluster membership >10% with a different species.

### 2.5. Population structure

Population structure in *C. dentata* was estimated using ADMIXTURE and a Discriminant Analysis of Principal Components (DAPC) (30). ADMIXTURE uses a model-based approach, like STRUCTURE, to estimate ancestry, but uses maximum likelihood rather than MCMC and is more computationally efficient (29). DAPC combines PCA and a Discriminant Analysis (DA) to identify demographically independent clusters by minimizing within group variance and maximizing between group variances (30). The VCF file was first converted to a BED file and linkage-disequilibrium (LD) pruned to include SNPs with R2 values < 0.1 within 50 SNP sliding windows (step size 10 SNPs) in PLINK v1.9 (31). ADMIXTURE was performed using the pruned BED file for K values 1-9. The –cv and -j120 options were enabled to allow for a five-fold cross validation and for the analysis to run in multithreaded mode using 120 threads, respectively. The lowest CV error score was used to determine the most likely value of K. The populations identified by ADMIXTURE were used for all subsequent analyses unless otherwise noted. Population membership for each sample was determined by highest ancestry proportion from the ADMIXTURE results.

The DAPC analysis was performed in R using the *adegenet* package (32). To create the input file for DAPC, the pruned BED file was converted back to a VCF file in PLINK v1.9 using the –recode vcf option and the –ref-from-fa option. The pruned VCF file was converted to genlight format using the vcfR package (33). The optimal number clusters was determined with the find.clusters function in *adegenet*, and the Bayesian Information Criterion (BIC) statistic was calculated for a maximum of five clusters and 355 principal components (PCs) (PCs = n-1, where n is the number of samples). The cluster number with the lowest BIC score was assumed to be the most likely value. The optim.a.score was assessed for each DAPC run to determine the optimal number of PCs to retain.

### 2.6. Demographic history

The sequential Markovian coalescent approach in SMC++ was used to estimate the historical effective population sizes (N_e_) of *C. dentata* from whole genome data (34). SMC++ required the input VCF file to not have any filtering for LD or MAF, so the input VCF file of 356 *C. dentata* was only filtered for high missing sites (>10% missingness) and biallelic SNPs. Sites missing in the *C. dentata* reference genome, which could be erroneously interpreted as long runs of homozygosity, were masked with a bed file generated with a conversion script (https://www.danielecook.com/generate-a-bedfile-of-masked-ranges-a-fasta-file/). The SMC files were generated for each of the 12 chromosomes using the vcf2smc function in SMC++. The estimate function was then used with a mutation per generation rate set to 5.2 × 10^−8^ from estimates of pedunculate oak (*Quercus robur*) (35). A 30-year generation time was assumed to convert coalescent events to years. A CSV file of the results was generated with the plot function and -c option for plotting in R (36).

### 2.7. Migration rates

Migration rates for *C. dentata* were estimated using Estimated Effective Migration Surfaces (EEMS) (37). EEMS estimates and visualizes effective migration and diversity across a given geographic range using genetic data from known locations (37). A stepping stone model is assumed and that isolation-by-distance is a component of the populations. The EEMS program requires three input files: an average pairwise differences matrix (DIFFS), list of sample geographic locations (COORD), and a list of habitat boundary coordinates (OUTER). To reduce computational load, the LD pruned *C. dentata* dataset from the population structure analyses was used. The pruned BED file was converted to the EEMS DIFFS file using the EEMS program bed2diffs. The habitat boundary coordinates were mapped with an online tool (http://www.birdtheme.org/useful/v3tool.html).

An initial run was performed with 8 million iterations, 1 million burn-in, 600 nDemes, 9999 thinning, and the hyperparameters in their default setting. EEMS relies on properly adjusted proposal variances, which influence the predicted migration and diversity rates. During the analysis, each of the output proposal acceptance frequencies should be between 20%-30%, but it is sufficient for them to range from 10%-40% (https://github.com/dipetkov/eems/blob/master/Documentation/EEMS-doc.pdf). Following the initial EEMS run, there were two proposal frequencies that were either less than 10% or greater than 40%. To account for this, the mEffectProposalS2 parameter was increased to 0.2 and mrateMuProposalS2 was decreased to 0.002. Four values of demes, 200, 350, 500, and 650 were used to evaluate the number of demes for best fit. Three chains, each with 20 million iterations, four-million burn-in, and 9999 thinning were performed for each deme and their outputs were combined in R using the rEEMSplots2 program. The posterior trace plot was evaluated to determine if the MCMC chain converged. Any EEMS run that did not converge was restarted from the final parameter state in the previous run. The results were mapped in R using rEEMSplots2. A linear regression line was fitted to the default ‘Dissimilarities between pairs of sampled demes’ plot for each deme value, and the deme value with the highest R^2^ value was determined to be the best fit. Appalachian Mountain peak locations were obtained from Wikipedia (https://en.wikipedia.org/wiki/List_of_mountains_of_the_Appalachians) and a smoothed loess line of these locations was added to the EEMS figure using *ggplot2*.

### 2.8. Tests for neutrality and nucleotide diversity

We used ANGSD to perform the population statistical analyses since its use of genotype likelihoods has been found to provide less biased estimates than previous methods that require genotype calling (38, 39). The sorted bam files generated from the SNP calling methods were used as the initial input for ANGSD to generate the SAF file (site allele frequency likelihood) for each population. Due to computational considerations, a SAF file was generated for each chromosome within a population using -doSaf 1 and then merged using realSFS cat. The filtering parameters used for each SAF file were adapted from ANGSD recommendations and other hardwood tree studies (40, 41). These parameters were: adjust mapping quality for excessive mismatches (-C 50), minimum base quality score 20 (-minQ 20), minimum mapping quality 30 (-minMapQ 30), discard reads that do not uniquely map (-uniqueOnly 1), only retain sites where the pair could be mapped (-only_proper_pairs 1), and remove ‘bad’ reads (-remove_bads 1). Additionally, to polarize the alleles in the site-frequency-spectrum (SFS), an ancestral reference fasta was generated in ANGSD using 11 *C. mollissima* BAM files with the following parameters: -doFasta 2 and -doCounts 1. Once the individual chromosomes were merged to generate a SAF file for each population, the SFS was calculated using realSFS.

Population statistics were computed for nucleotide diversity and tests of neutrality in ANGSD. Thetas were first calculated from the SFS in ANGSD using realSFS saf2theta. The statistics for each chromosome were then determined using thetaStat do_stat. Nucleotide diversity was calculated from the thetaStat do_stat output by dividing the pairwise theta (tP) by the number of sites (-nSites) evaluated by ANGSD for that genomic region. A sliding window analysis was also performed on each chromosome using a 50 kb window and a 10 kb slide.

Pairwise F_ST_ was evaluated using ANGSD for each of the populations to estimate the population differentiation of *C. dentata* and identify candidate regions of the genome undergoing selection. Using the per chromosome SAF files as input, each pair of populations per chromosome were used to generate separate pairwise 2D-SFS files. The F_ST_ index was then performed on each pairwise 2D-SFS file using realSFS fst index. A sliding window analysis was then performed for the F_ST_ calculation with a 50 kb window and a 10 kb slide. The estimated pairwise F_ST_ for each chromosome was averaged for all 12 chromosomes to get the mean pairwise F_ST_ between populations.

VCFtools was used to calculate the observed heterozygosity for each sample. The filtered SNP dataset was used as input with the options -s – and -het. The output file provides the observed and expected homozygosity, F statistic, and number of nucleotide sites analyzed for each sample. To calculate the observed heterozygosity for each sample, we first subtracted the observed homozygosity from the number of sites, and then divided that total from the number of sites. This provides the ratio of observed heterozygosity for each sample. To calculate the average observed heterozygosity for each population, the sample memberships from the ADMIXTURE analysis were used. The observed heterozygosity was averaged over all samples within a population to calculate the average observed heterozygosity for that population. To determine the expected heterozygosity for each sample and each population, the previous steps were performed using the expected homozygosity values in place of observed homozygosity. Observed heterozygosity between all populations was tested for significance using a one-way ANOVA test in R with function aov(). Further comparisons between each pair of populations were completed using Tukey multiple pairwisecomparisons with R function TukeyHSD(). For the one-way ANOVA and Tukey tests, a P-value of 0.05 was used for significance.

### 2.9. Detecting signatures of positive selection

We used RAiSD to identify genomic regions undergoing positive selection in each population (42). RAiSD estimates a composite statistic, µ, which evaluates each genomic region based on multiple neutrality and diversity metrics. RAiSD has been shown to excel at identifying regions that are undergoing selective sweeps, while being more computationally efficient than other leading methods (42). The filtered *C. dentata* VCF file was subset for each of the three populations identified by ADMIXTURE and used as input. Regions that were missing data in the *C. dentata* reference genome, denoted with N, were masked with the -X flag and a BED file of missing locations. The default parameters were used for each run, except that we assigned a seed for the random number generator. The RAiSD filtering parameters for each population’s dataset retained 8,833,026 SNPs for the northeast population, 7,545,294 SNPs for the central population, and 11,737,005 SNPs for the southwest population for analysis. We used the quantile function in R (36) to identify the 0.1% outlier regions in each of the RAiSD site reports for each population. Results of this analysis were displayed using *ggplot2* (43).

To identify the genes associated with the 0.1% outlier regions for each population, we obtained gene names and location information from the American chestnut genome feature file (Cdentata_673_v1.1.gene.gff3.gz; *Castanea dentata* v1.1; http://phytozome-next.jgi.doe.gov/). Genes were determined to be associated with the RAiSD outlier regions if they resided within 1 Kb of the region. Only unique genes were retained. To determine gene function, we identified the orthologs for each of the outlier genes in *Arabidopsis thaliana*. The list of outlier *C. dentata* genes for each population were entered into BioMart on Phytozome (44, 45), and the *A. thaliana* TAIR10 genome was used to output a list of corresponding orthologs (46). The list of *A. thaliana* orthologs were entered into the online TAIR gene ontology tool to obtain the function category for each ortholog (47)(https://www.arabidopsis.org/tools/bulk/go/index.jsp). Finally, we performed GO enrichment analyses for biological function on four gene sets. These sets were the unique genes belonging to the southwest, central, and northeast populations, and the set of genes that are shared between all three populations. The *A. thaliana* orthologs were retrieved for each gene set using the previously described methods, and were submitted to the TAIR GO Term Enrichment for Plants tool (https://www.arabidopsis.org/tools/go_term_enrichment.jsp), which sends the data to the PANTHER Classification System (48). The summary for each gene set analysis using the PANTHER Classification System was as follows. Analysis type: PANTHER Overrepresentation Test (Released 20210224), Annotation Version and Release Date: GO Ontology database DOI: 10.5281/zenodo.5228828 Released 2021-08-18, Reference list: *Arabidopsis thaliana* (all genes in database), Annotation Data Set: GO biological process complete, Test Type: Fisher’s Exact, Correction: Bonferonni correction for multiple testing for p<0.05.

## 3. Results

### 3.1. Genomic datasets

Of the 384 samples sequenced, 86 had greater than 20x coverage, 242 had 10-20x coverage, and 56 had less than 10x coverage. The 384 sample *C. dentata* VCF file contained 23,720,251 SNPs and INDELs. Eighteen samples with greater than 10% missing data were removed, and 10 additional samples were removed that had > 10% cluster membership with one or more of the *Castanea* species reference samples in ADMIXTURE analysis. The final *C. dentata* dataset contained 356 individuals with an average coverage of ≈ 17x and 21,136,994 high quality SNPs (Fig. S1). The pruned *C. dentata* dataset contained 3,539,550 SNPs. The *Castanea* species reference dataset contained 92 samples and 62,647,079 SNPs that passed the filtering criteria. The combined and filtered *C. dentata* and *Castanea* species reference dataset contained 76,378,648 SNPs and INDELs.

### 3.2. Hybridization

ADMIXTURE analysis with the combined *C. dentata* and *Castanea* species reference panel suggested seven clusters best explained the data, with each *Castanea* species as an individual cluster in addition to three clusters within *C. dentata* (Fig. S2). Of the 384 putative American chestnut individuals, 340 had >99% ancestry assigned to the *C. dentata* clusters. However, ten individuals showed a significant level of ancestry from other *Castanea* species (>10%) and were removed from further analyses. Three samples were identified as *C. pumila*, four samples were *C. sativa* x *C. dentata* hybrids, and three were *C. mollissima* x *C. dentata* hybrids. Overall, the American chestnut samples sequenced did not reveal widespread patterns of significant introgression with other *Castanea* species.

### 3.3. Population structure in *Castanea dentata*

Population structure within *C. dentata* was best explained by a two or three population model as identified by the DAPC BIC plot and ADMIXTURE CV error plot, respectively (Fig. 2). The three population ADMIXTURE model was characterized by a southwest, central, and northeast cluster (Fig. 2). The southwest and central population separated in northern Georgia and eastern Tennessee, while the central and northeast population have an area of admixture in Pennsylvania before becoming more distinctly separated in southern New York. The two population DAPC model included the same southern population and boundary as ADMIXTURE, but the central and northeastern populations were merged. Both analyses were mostly in agreement with population memberships at the same K values.

**Fig. 2.**
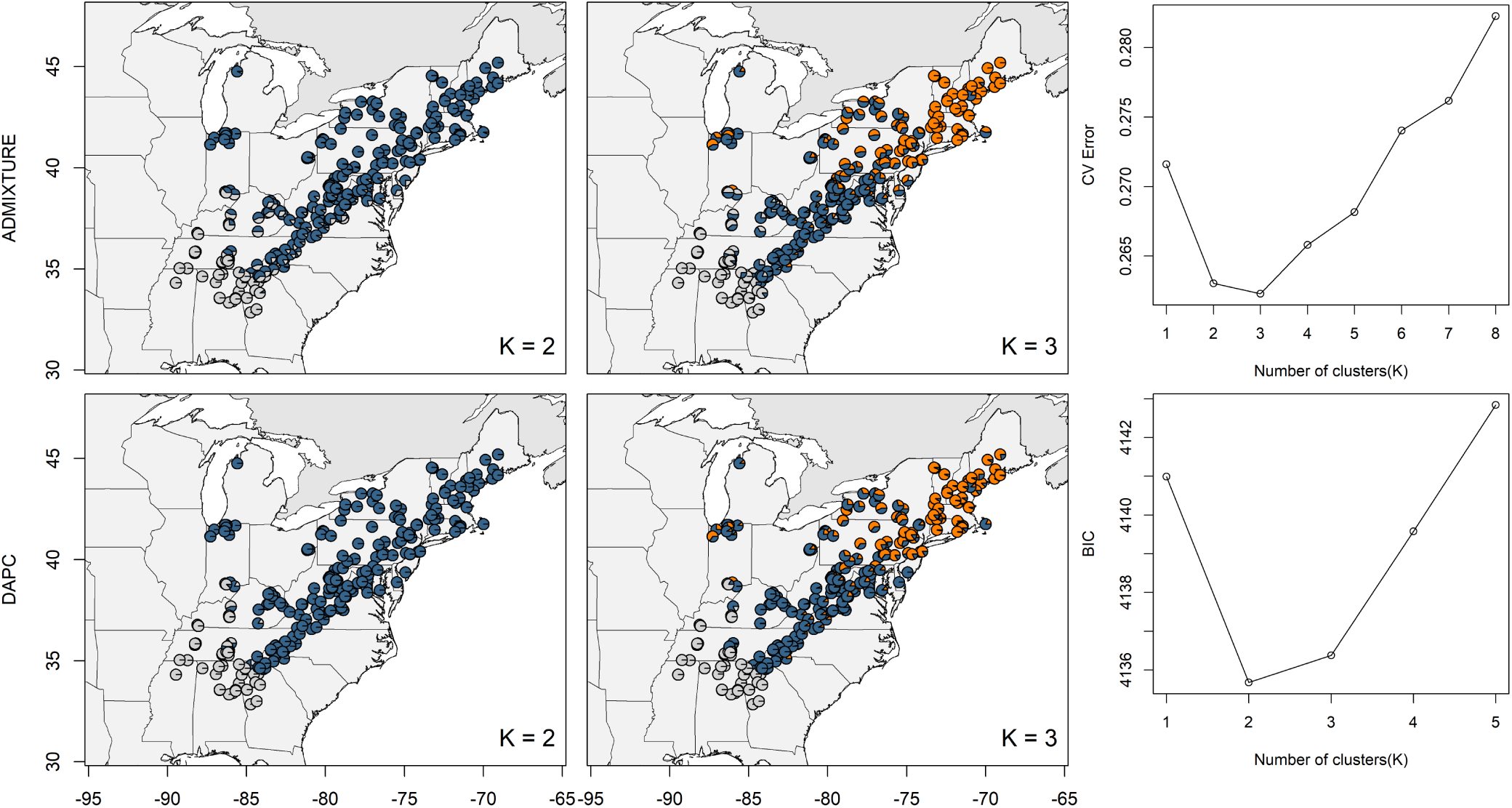
The wild American chestnut population structure is best described by three or two populations. The pie charts represent the sample locations and the proportion of color within each pie chart represents population membership. The minimum CV error score and minimum BIC score for each K value (right) were used to determine most likely population number for the ADMIXTURE and DAPC analyses, respectively.

### 3.4. Migration rates

The R^2^ value for the dissimilarity plots for the 650 deme model (R^2^ = 0.92) and 500 deme model (R^2^ = 0.93) were similar, and we present the 650 deme model due its increased resolution. EEMS analysis suggests that the Appalachian Mountains form a barrier to gene flow along their length, with migration running from southeast to northeast on either side of the mountain range (Fig. 3a). A single region of above average effective migration rate was shown in southern West Virginia that crosses the Appalachian Mountain range (Fig. 3a). The ADMIXTURE K=4 model agrees with the EEMS estimates and suggests a further subdivision of the central population on either side of the Appalachian Mountains (Fig. 3b). EEMS also estimates a diversity parameter (q), which reflects genetic dissimilarity between individuals within the same deme, and can thus be thought of as the within-population component of genetic variance. This diversity parameter was generally high throughout the range, though more so in the central and southern portion of the range, and somewhat lower in the northeast and northwest (Fig. S3). Pockets of lower diversity, and thus higher intra-deme genetic similarity, on the outer edges of the species native range may reflect areas of more recent expansion-associated bottlenecks (Fig. S3).

**Fig. 3.**
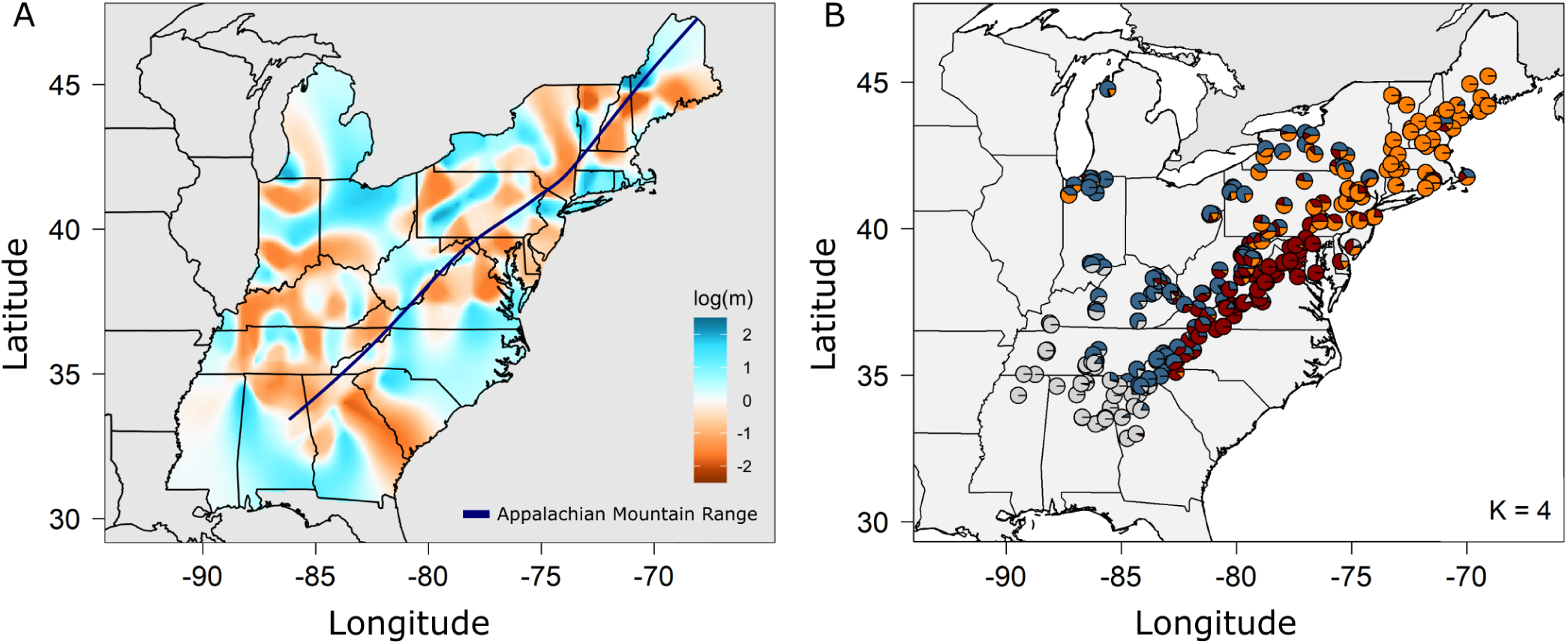
The Appalachian Mountain range was a barrier to gene flow for postglacial migrating American chestnut populations. **A**. EEMS migration rate estimates for American chestnut populations. Regions of blue are above average migration estimates, while orange regions are below average migration estimates. A loess line of Appalachian Mountain peaks was applied in *ggplot2* using peak locations obtained from https://en.wikipedia.org/wiki/List_of_mountains_of_the_Appalachians. **B**. ADMIXTURE analysis with K = 4. The pie charts represent sample location and the color represents population membership.

### 3.5. Demographic history

SMC++ estimates of N_e_ over time suggest that each population underwent contractions and expansions in N_e_, beginning approximately two million years ago. All populations followed a similar pattern of demographic history, however, the southwest population lagged the central and northeastern populations’ events. N_e_ rapidly increased for all three populations approximately 6,700-11,700 years ago, after which the central population underwent an additional contraction within the past 7,000 years (Fig. 4). The southwest population had the highest contemporary N_e_ (N_e(southwest)_=20,306, N_e(central)_= 8,347, N_e(northeast)_= 13,078).

**Fig. 4.**
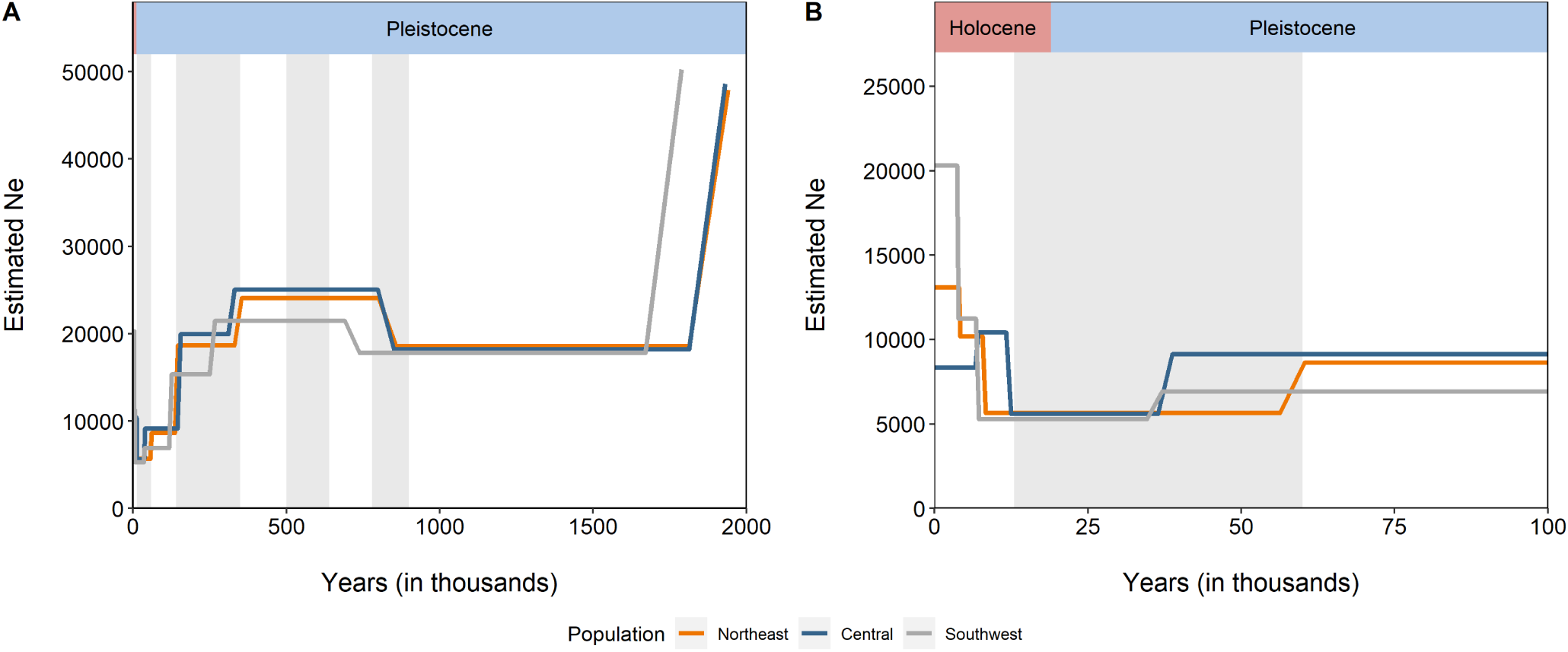
American chestnut populations underwent bottleneck and expansion events that coincide with recent North American glaciations. Greyed regions represent glaciation event times approximated from Bowen (49) (as cited in (50)). Each colored line represents the N_e_ for each of the three populations. Coalescent scaling was converted to years with a 30 year generation time. **A**. SMC++ N_e_ estimates from the present to 2 million years ago. **B**. SMC++ N_e_ estimates from the present to 100,000 years ago.

### 3.6. Genomic diversity and tests of neutrality

The southwest population had the greatest nucleotide diversity, followed by the central and northeast populations (π_southwest_ = 0.0069; π_central_ = 0.0064; π_northeast_ = 0.0058). All populations had negative average Tajima’s D, which were similarly clinal(D_southwest_= -1.083; D_central_=-1.016; D_northeast_=- 0.335). Consistent with these negative values for Tajima’s D, the SFS plot revealed that for each population, there was a deficiency of rare variants and an excess of high frequency variants (Fig. S4). Sliding window analyses revealed heterogeneous genome-wide Tajima’s D, nucleotide diversity, and F_ST_ (Fig. 5). Throughout the genome, the southwest population had the most negative Tajima’s D values, followed by the central and northeast populations (Fig. 5). Conversely, the southwest population had the highest nucleotide diversity values throughout the genome, with decreasing values for the central and northeast populations (Fig. 5). The highest F_ST_ values were attributed to the southwest-northeast population pair (Fig. 5).

**Fig. 5.**
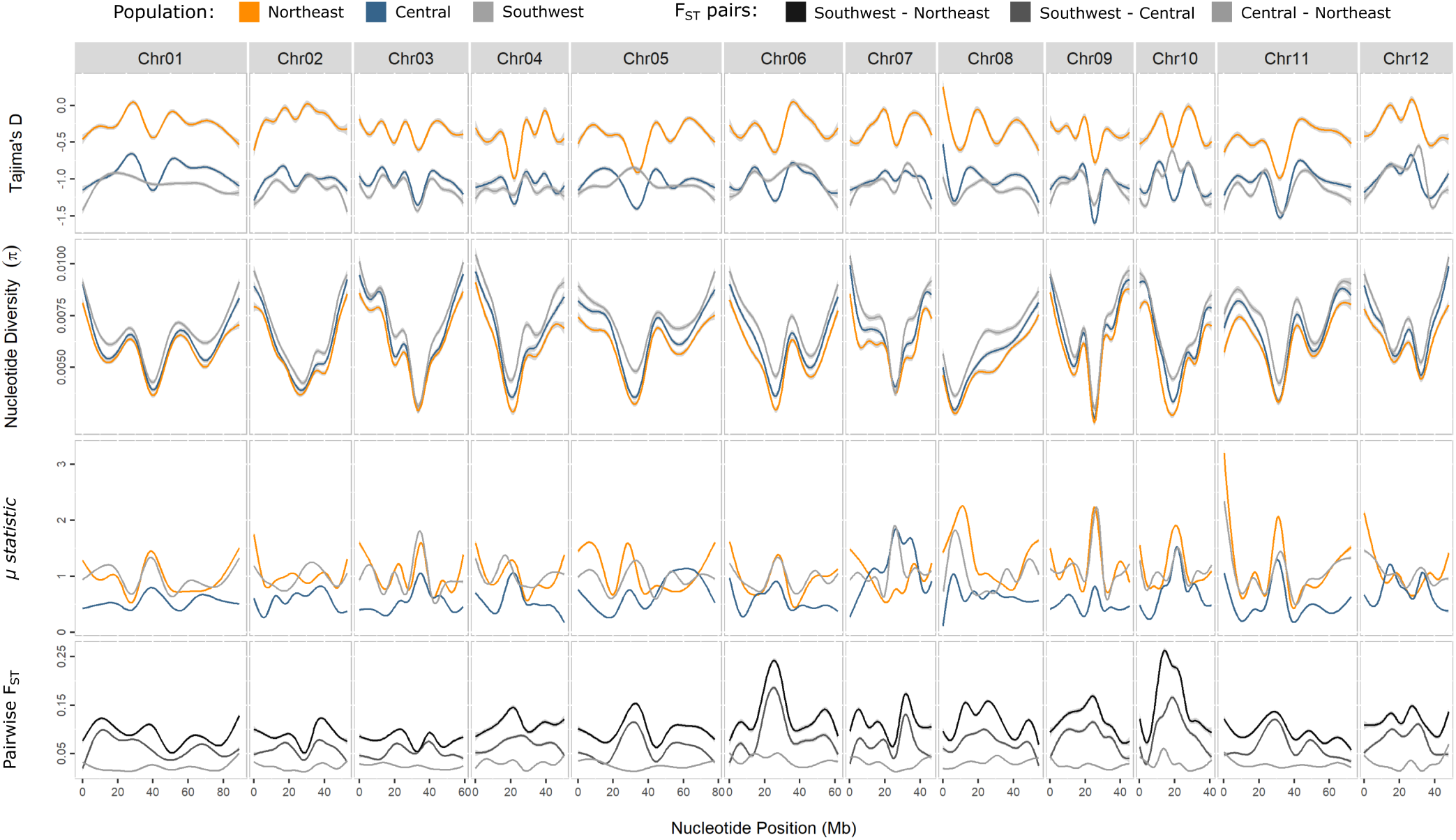
Genome-wide sliding-window analysis of the three American chestnut populations for Tajima’s D, nucleotide diversity (π), RAiSD µ-statistic, and pairwise F_ST_. Smoothed lines were computed for all results in *ggplot2* with the default gam parameter.

Consistent with the pattern for nucleotide diversity, heterozygosity was highest in the southwest (Table 1). Northeast H_o_ was significantly lower than the central and the southwest population (p<0.001, p<0.001) (Fig. S5a). F_ST_ estimates between population pairs were relatively low for all comparisons, with the highest divergence between the southwest and northeast populations (F_ST(southwest-northeast)_ =0.1076, F_ST(southwest-central)_ = 0.0705, F_ST(central-northeast)_ = 0.0268).

**Table 1.** Average Tajima’s D, nucleotide diversity, heterozygosity, and inbreeding coefficient for each American chestnut population.

| Population | Tajima's D | $\pi$ | $H_o$ | $H_e$ | F |
| --- | --- | --- | --- | --- | --- |
| Southwest | -1.0827 | 0.0069 | 0.1515 | 0.1660 | 0.0882 |
| Central | -1.0156 | 0.0064 | 0.1492 | 0.1660 | 0.1016 |
| Northeast | -0.3345 | 0.0058 | 0.1353 | 0.1660 | 0.1851 |

### 3.7. Genomic regions and associated genes undergoing positive selection

Every chromosome for each population contained outlier genomic regions identified by RAiSD (Fig. 5). The southwest population had the greatest number of significant regions with 11,733, followed by the northeast population with 8,832, and the central population with 7,534. Within the significant regions, the southwestern population contained 617 outlier genes, which was the most out of the three populations (Fig. 6). Among these outlier genes, 402, 387, and 323 were unique to the southwestern, central, and northeastern populations, respectively, while 49 genes were shared between all three populations (Fig. 6).

**Fig. 6.**
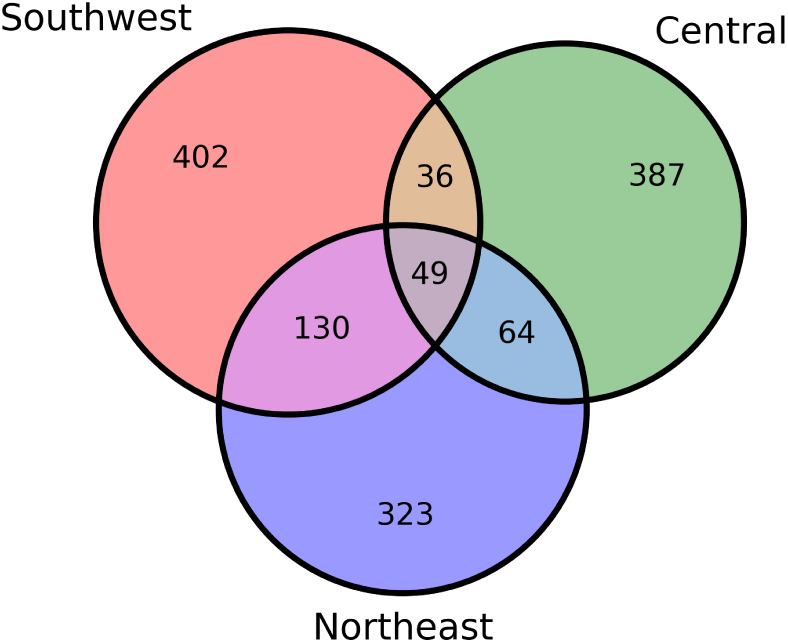
Number of outlier genes identified by RAiSD within each American chestnut population.

### 3.8. GO enrichment analysis

GO enrichment analysis revealed that “response to stress” was within the top four most annotated gene families for the GO biological process for each population (Fig. S6). Among the unique genes within each population, the southwest population had the most overrepresented GO terms, with 25 (Table S1), while the northeast population had 10 (Table S2), and the central population contained one (Table S3). The GO term “defense response to fungus” was significantly overrepresented in the shared gene set for all three populations (Fold enrichment = 16.5, p = 0.0373). The overrepresented genes for the shared gene set were Caden.02G006600 and Caden.03G057600, which are both in the apoptotic ATPase eukaryotic orthologous group (KOG).

## 4 Discussion

The American chestnut was an economically and ecologically important tree species that was decimated by an invasive fungal blight approximately 100 years ago. Blight resistant American chestnut populations are being developed, and these populations will need sufficient genomic diversity to thrive across the diverse and rapidly changing climatic gradient of the species historical range. Our goal is to prioritize areas for *ex situ* conservation through propagation of wild trees. These trees will then be used to introgress adaptive genetic variation into chestnut blight-resistant populations. The first step in this process is to define broad management units on the basis of population structure and postglacial history for the species (13).

To address this goal, we re-sequenced 384 wild chestnut genotypes, which yielded over 23 million variants. These data revealed that population structure in chestnut can be best described by a two or three population model, with a southwestern population being present in both models. The southwestern population was the most genetically diverse, and that diversity decreases as latitude increases. These contemporary patterns of genomic diversity in American chestnut are most likely the result of past population size reductions, and recent expansions, associated with Quaternary glaciation of North America.

### 4.1. Identification of hybridization in American chestnut

Our primary goal in developing a WGS dataset that included most *Castanea* species was to exclude hybrids between our wild *C. dentata* samples and congeners a significant concern given the history of planting and naturalization of non-native *Castanea* species in the eastern U.S. (51). We did not detect widespread introgression with other chestnut species, which is consistent with previous assessments of North American *Castanea* (17). However, our *C. dentata* sampling focused on trees with morphological characteristics representative of the species. Thus, our sample was biased against genotypes displaying intermediate phenotypes, and hybrids with naturalized non-native *Castanea* species, as well as the native *C. pumila*, may nevertheless be present in the wild. A systematic exploration of the relationships among worldwide *Castanea* species is beyond the scope of this paper, and future studies will use these data to resolve the phylogenetic relationships between *Castanea* species and within the *Castanea pumila* species complex.

### 4.2. Population structure

Our range-wide sample of 356 American chestnut genotypes suggests either two or three genetically distinct populations, separated along a north-south gradient. Both the DAPC and ADMIXTURE analyses agreed with the presence of a southwestern population, but differed on the number of populations in the remainder of the range. Nevertheless, both DAPC and ADMIXTURE gave very similar cluster memberships and patterns at each K value. Previous studies primarily used microsatellite data to describe the population structure and genetic diversity of the American chestnut. Kubisiak and Roberds (14) sampled several locations from North Carolina through Massachusetts and found that a single, genetically diverse metapopulation best fit the data. Two additional studies used sampling sites from Kubisiak and Roberds (14) but analyzed a different set of microsatellite markers. Gailing and Nelson (15) and Müller et al. (16) found that the data best supported two populations — a northeastern and southwestern population – with Pennsylvania and Maryland being a transition zone between these two groups. These studies did not sample south of North Carolina, which may explain their two population estimates. More recently, Spriggs and Fertakos (17) used ddRAD-seq data and identified the southern portion of the range as a separate population after a more extensive sampling of the southern region. Their results agreed with our DAPC analysis, whereas our ADMIXTURE analysis suggests a further division between the central and northern portions of the range. This may be due to the increased genomic coverage of our study, our larger sample size and more uniform range-wide sampling, or some combination of these factors.

While the scarcity of viable seed produced by wild American chestnuts make common gardens difficult, one such study of 13 American chestnut provenances, collected from throughout the native range *C. dentata* and planted in Vermont, found that seedlings from warmer and moderate climates grew faster but had greater winter injury than northern seed sources (52–54). The same study showed that nuts collected from the north have greater winter cold hardiness than those from the south (53). Thus, like most temperate and boreal tree species, American chestnut exhibits local adaptation to climate. The three populations we identified appear to be differentiated at latitudinal temperature and precipitation breaks along the Appalachian Mountain range. Pennsylvania marks the region of admixture between the central and northeastern populations, and is also where mean winter temperatures transition from above freezing to below freezing (2015 PRISM climate group, https://prism.oregonstate.edu/normals/). In addition, the separation between the central and the southwestern populations occurs in Tennessee, which is the northern boundary for an area of the southern U.S. that receives 250mm higher annual precipitation on average compared with eastern and northern portions of the historical chestnut range (https://prism.oregonstate.edu/normals/). These three populations may thus comprise broad chestnut ecoregions, reflecting both isolation-by-distance but also potentially isolation-by-adaptation (55–57).

### 4.3. Patterns of genomic diversity

Genome-wide nucleotide diversity in American chestnut was comparable to other widely distributed forest tree species (58). The southern portion of the range had the highest levels of nucleotide diversity, which decreased as latitude increased, consistent with previous studies (14–17). Mean heterozygosity estimates across all populations were much lower, and the inbreeding coefficient higher, than previous estimates from microsatellites (15). F_ST_ between populations was relatively low, with the greatest divergence between the southwest and northeast populations. Tajima’s D was negative in each population, which was driven by an excess of rare variants for each population.

The combination of negative Tajima’s D, low levels of heterozygosity, and an excess of rare variants suggest that all populations are undergoing expansion following recolonization bottlenecks, the timing of which varied by population. The more negative values of Tajima’s D for the southwest population, followed by the central and northeastern populations, suggest that the southwest population underwent a more ancient bottleneck. For nucleotide diversity, we observed the inverse. The southwest population has the highest levels of nucleotide diversity, which decreased as latitude increases, suggesting that the southwestern population has a higher long term effective population size, and was most likely a glacial refugium from which postglacial expansion occurred. This same pattern of inverse relationships between neutrality and diversity tests can be found in Sitka spruce (*Picea sitchensis*), which underwent recolonization of its northern range from a southern glacial refugium (59). The patterns in genome-wide diversity estimates among populations may also be due to different pressures of natural selection (60). Additionally, we found that the more geographically separated populations were more genetically divergent, which is similar to other temperate species (40, 61). Though, the overall genetic distinctness between populations was relatively low, with most of the genomic diversity being accounted for within populations findings that are consistent with a multi-species assessment of northeastern North American tree species (62).

### 4.4. Demographic history and migration patterns

N_e_ for each of our populations declined in the distant past and subsequently each experienced repeated size changes before a rapid expansion within the past 11,000 years. In the distant past, changes in N_e_ for the northeastern population generally paralleled those of the central population, while the southwest population lagged, likely due to more muted impacts of climate change in this area. Curiously, following an initial increase after the last glacial maximum, N_e_ for the central population size again decreased in the recent past. This decline may be due to a recent bottleneck event, or uncertainty in the SMC++ analysis, however, it was not due to the chestnut blight. The decline of the American chestnut populations from the blight occurred within the last century, and most of the trees we sampled are stump sprouts derived from surviving root stock that predate the blight. Thus, the recent dramatic reduction of American chestnut census population size would not influence our demographic history analyses.

The population declines and subsequent expansion follow the Quaternary glaciation events, which began approximately 2.7 million years ago (63). Previous biogeographical assessments for the eastern United States (64), as well as genetic studies, suggest that tree species in eastern North America migrated from southwestern refugia (15, 17). Fossil pollen evidence indicates the Gulf coastal forests in Florida and Southern Alabama were a glacial refuge for *C. dentata* during the Wisconsonian glaciation approximately 25,000 – 31,000 year ago (51, 65, 66). As the glaciers retreated, pollen evidence indicates that *C. dentata* migrated north into Tennessee approximately 15,000 years ago (67, 68), and continued northeastward along the Appalachian Mountains (68) at a rate of approximately 100 meters per year, eventually arriving in Connecticut approximately 2,000 years ago (64). However, other glacial refugia may have existed more northeastward and the rate of migration northward may have been slower than what fossil pollen suggests (69). As populations migrated northeastward and diverged, bottlenecks likely occurred, leading to a northeastern population with less genomic diversity than those further south.

When we allowed for four clusters in the ADMIXTURE analysis, the central population separates on either side of the Appalachian Mountain range. The Appalachian Mountain range may thus serve as a barrier to gene flow throughout the American chestnut range, with migration paths northeastward on either side. With four clusters, we observed a region of mixing between eastern and western clusters in southwest Virginia and southeast West Virginia, which the EEMS analysis showed was also an area of enhanced gene flow. A migration route may have existed in this area, which may explain the observation by Gailing and Nelson (15) that chestnuts in Ontario, Canada were more similar to the North Carolina population than their northeastern neighbors.

### 4.5. Signatures of selection in American chestnut populations

The ANGSD sliding window analyses revealed several broad genomic regions of negative Tajima’s D and reduced nucleotide diversity. Some of these regions likely reflect reduced recombination and associated decreased nucleotide diversity near centromeres. However, several intervals, such as on chromosome six and chromosome ten, also had elevated F_ST_ for the southwest-central and southwest-northeast population pairs, suggesting that these regions may have experienced recent selection related to environmental adaptation to more northern climates. RAiSD identified several thousand outlier regions that may be targets of selection that were enriched for genes related to “response to stress” and “response to chemical”. Further, the GO enrichment analysis identified “defense response to fungus” as the only overrepresented biological pathway for the set of shared genes between all populations. This suggests that selection related to pathogen pressure is a key feature of global adaptation in American chestnut.

A few of the American chestnuts sampled in this study were large surviving trees that may have low levels of blight resistance, however, most were blight killed resprouts. Furthermore, American chestnuts from across the species range are highly susceptible to *Phytophthora cinnamomi* (the Oomycete responsible for *Phytophthora* root rot) - a contemporary agent of American chestnut decline (70, 71). To date, the only known source of *P. cinnamomi* resistance for *C. dentata* has been introgression from Asian *Castanea* species, such as *C. mollissima* and *C. crenata* (71–73). Thus, the enrichment for defense genes among selection targets we observed is unlikely to be related to these two contemporary pathogens that threaten the species. Rather, it likely reflects historical interactions with, and adaptations to, as yet unknown native fungal pathogens.

This is not to say, however, that the “defense response to fungus” selection targets could have no role in responding to chestnut blight or *Phytophthora* root rot. In response to blight inoculation, both resistant *C. mollissima* and susceptible *C. dentata* show transcriptional responses, though more genes potentially related to blight resistance were upregulated in *C. mollissima* (74). Additionally, blight resistance in American chestnut backcross populations is polygenetic, with multiple loci contributing to resistance (75). The lack of blight resistance in American chestnut may be due to an inadequate or inappropriate transcriptional response to infection. As such, further evaluation of these genes is necessary to determine the cause of their overrepresentation and their possible relationship to blight resistance in wild populations.

### 4.6. Conclusion

We developed two high quality WGS datasets that will further population genomics studies of American chestnut and other *Castanea* species, which revealed that American chestnut underwent a postglacial migration northward that most likely influenced its current genetic structure. Three populations were identified that were separated along a latitudinal gradient, with the southern population having the highest levels of genetic diversity, which suggests it is most likely the oldest population and the refugia from which postglacial expansion occurred. Subtle population structure also revealed a separation of the central population on either side of the Appalachian Mountains that suggests these mountains represent a barrier to gene flow. We identified genomic targets of selection that were both unique to each population, and shared among all populations, which reflect adaptation to both the abiotic and biotic environments. Future breeding and conservation plans will need to consider these separate populations to preserve unique areas of American chestnut genetic diversity.

While this study describes the patterns of genetic structure that exist within wild American chestnut, it stops short of identifying the genomic signatures of local climate adaptation within each population. Failure to account for traits related to climate within breeding populations could lead to the reintroduction of maladapted blight-resistant trees that fail to compete with other native species (76). Identifying the genomic targets of climate-related selection is a key step in understanding the underlying genomic basis of local adaptation. Our future goal is to characterize the genomic architecture of local adaptation across the species range, and combine those results with this study’s findings to develop strategies for germplasm conservation and breeding to explicitly account for local adaptation in restoration populations.

## ACKNOWLEDGEMENTS

We would like to thank TACF volunteers who participated in sampling of wild trees for this project, as well as Dr. Rita Costa (Instituto Nacional de Investigação Agrária e Veterinária, Portugal) for providing *Castanea sativa* samples. We thank the HudsonAlpha Institute for Biotechnology and TACF for prepublication use of the genome of *Castanea dentata* funded by the Colcom Foundation. We thank Advanced Research Computing at Virginia Tech for providing computational resources, Dr. Robert Settlage for technical support related to the analyses described here. This research was supported by the United States National Institute for Food and Agriculture Projects 1018599 and 1005394, and by a Graduate Fellowship to AS from the Virginia Tech Institute for Critical Technology and Applied Science.

## DATA ACCESSIBILITY AND BENEFIT-SHARING SECTION

Data Accessibility Statement

### Genomic Data

The genomic sequences reported in this paper have been deposited in the NCBI SRA (BioProject PRJNA804196)

### Sample metadata

Metadata are also stored in the SRA (BioProject PRJNA804196)

### Scripts

The scripts used for the SNP calling pipeline and other mentioned scripts can be found at https://github.com/alex-sandercock/American_chestnut_WGS

### Benefit-Sharing Statement

Benefits Generated: Benefits from this research include the sharing of our genomic data as listed in the Data Accessibility Statement.

## AUTHOR CONTRIBUTIONS

A.M.S., J.G., J.A.H., J.S., and J.W.W. designed the study. Q.Z., H.A.J., T.M.S., J.A.S., S.F.F., K.C., J.S., and J.G. contributed to data collection. A.M.S. analyzed the data. A.M.S., J.A.H., and J.W.W. wrote the manuscript with input from all authors.

## References

1. G Geoff Wang, Benjamin O Knapp, Stacy L Clark, and Bryan T Mudder. The silvics of Castanea dentata (marsh.) borkh., american chestnut, fagaceae (beech family). In: Publication: Gen. Tech. Rep. SRS-GTR-173. Asheville, NC: US Department of Agriculture Forest Service, Southern Research Station, page 18, 2013.

2. Emily WB Russell. Pre-blight distribution of Castanea dentata (marsh.) borkh. Bulletin of the Torrey Botanical Club, 114(2):183–190, 1987.

3. Philip Laurence Buttrick. Chestnut and the chestnut blight in North Carolina. Number 56. 1925.

4. Samuel Detwiler. The american chestnut tree: Identification and characteristics. American Forestry, 21(262):957–959, 1915.

5. Richard A Jaynes and Arthur H Graves. Connecticut hybrid chestnuts and their culture. Connecticut Agricultural Experiment Station, 1963.

6. David M Smith. American chestnut: Ill-fated monarch of the eastern hardwood forest. Journal of Forestry, 98(2):12–15, 2000.

7. Ralph H Lutts. Like manna from god: The american chestnut trade in southwestern virginia. Environmental History, 9(3):497–525, 2004.

8. John Scrivani. Forest inventory and analysis. Journal of the American Chestnut Foundation, 25:17–18, 2011.

9. Frederick L Paillet. Chestnut: history and ecology of a transformed species. Journal of Biogeography, 29(10-11):1517–1530, 2002.

10. Steven L Stephenson, Harold S Adams, and Michael L Lipford. The present distribution of chestnut in the upland forest communities of virginia. Bulletin of the Torrey Botanical Club, 118(1):24–32, 1991.

11. Sean Hoban, Michael W Bruford, W Chris Funk, Peter Galbusera, M Patrick Griffith, Catherine E Grueber, Myriam Heuertz, Margaret E Hunter, Christina Hvilsom, Belma Kalamujic Stroil, Francine Kershaw, Colin K Khoury, Linda Laikre, Margarida Lopes-Fernandes, Anna J MacDonald, Joachim Mergeay, Mariah Meek, Cinnamon Mittan, Tarek A Mukassabi, David O’Brien, Rob Ogden, Clarisse Palma-Silva, Uma Ramakrishnan, Gernot Segelbacher, Robyn E Shaw, Per Sjögren-Gulve, Nevena Veličković, and Cristiano Vernesi. Global commitments to conserving and monitoring genetic diversity are now necessary and feasible. BioScience, 71(9):964–976, 2021.

12. Jared W Westbrook, Jason A Holliday, Andrew E Newhouse, and William A Powell. A plan to diversify a transgenic blight-tolerant american chestnut population using citizen science. Plants, People, Planet, 2(1):84–95, 2020.

13. W Chris Funk, John K McKay, Paul A Hohenlohe, and Fred W Allendorf. Harnessing genomics for delineating conservation units. Trends in Ecology & Evolution, 27(9):489–496, 2012.

14. Thomas L Kubisiak and James H Roberds. Genetic structure of american chestnut populations based on neutral dna markers. In Restoration of American chestnut to forest lands: proceedings of a conference and workshop. Asheville, NC, USA: National Park Service, pages 109–122, 2006.

15. Oliver Gailing and C Dana Nelson. Genetic variation patterns of american chestnut populations at est-ssrs. Botany, 95(8):799–807, 2017.

16. Markus Müller, C Dana Nelson, and Oliver Gailing. Analysis of environment-marker associations in american chestnut. Forests, 9(11):695, 2018.

17. Elizabeth L Spriggs and Matthew E Fertakos. Evolution of Castanea in north america: restriction-site-associated dna sequencing and ecological modeling reveal a history of radiation, range shifts, and disease. American Journal of Botany, 108(9):1692–1704, 2021.

18. David B Lowry, Sean Hoban, Joanna L Kelley, Katie E Lotterhos, Laura K Reed, Michael F Antolin, and Andrew Storfer. Breaking rad: An evaluation of the utility of restriction site-associated dna sequencing for genome scans of adaptation. Molecular Ecology Resources, 17:142–152, 2017.

19. Peter Tiffin and Jeffrey Ross-Ibarra. Advances and limits of using population genetics to understand local adaptation. Trends in Ecology & Evolution, 29(12):673–680, 2014.

20. Badr Benjelloun, Frédéric Boyer, Ian Streeter, Wahid Zamani, Stefan Engelen, Adriana Alberti, Florian J Alberto, Mohamed BenBati, Mustapha Ibnelbachyr, Mouad Chentouf, Abdelmajid Bechchari, Hamid R Rezaei, Saeid Naderi, Alessandra Stella, Abdelkader Chikhi, Laura Clarke, James Kijas, Paul Flicek, Pierre Taberlet, and François Pompanon. An evaluation of sequencing coverage and genotyping strategies to assess neutral and adaptive diversity. Molecular Ecology Resources, 19(6):1497–1515, 2019.

21. Ellen Crocker, Bradford Condon, Abdullah Almsaeed, Benjamin Jarret, C Dana Nelson, Albert G Abbott, Doreen Main, and Margaret Staton. Treesnap: A citizen science app connecting tree enthusiasts and forest scientists. Plants, People, Planet, 2(1):47–52, 2020.

22. Geraldine A Van der Auwera and Brian D O’Connor. Genomics in the cloud: using Docker, GATK, and WDL in Terra. O’Reilly Media, 2020.

23. Heng Li and Richard Durbin. Fast and accurate long-read alignment with burrows–wheeler transform. Bioinformatics, 26(5):589–595, 2010.

24. Heng Li, Bob Handsaker, Alec Wysoker, Tim Fennell, Jue Ruan, Nils Homer, Gabor Marth, Goncalo Abecasis, and Richard Durbin. The sequence alignment/map format and samtools. Bioinformatics, 25(16):2078–2079, 2009.

25. Aaron McKenna, Matthew Hanna, Eric Banks, Andrey Sivachenko, Kristian Cibulskis, Andrew Kernytsky, Kiran Garimella, David Altshuler, Stacey Gabriel, Mark Daly, and Mark A DePristo. The genome analysis toolkit: a mapreduce framework for analyzing next-generation dna sequencing data. Genome Research, 20(9):1297–1303, 2010.

26. Ryan Poplin, Valentin Ruano-Rubio, Mark A DePristo, Tim J Fennell, Mauricio O Carneiro, Geraldine A Van der Auwera, David E Kling, Laura D Gauthier, Ami Levy-Moonshine, David Roazen, Khalid Shakir, Joel Thibault, Sheila Chandran, Chris Whelan, Monkol Lek, Stacey Gabriel, Mark J Daly, Ben Neale, Daniel G MacArthur, and Eric Banks. Scaling accurate genetic variant discovery to tens of thousands of samples. BioRxiv, page 201178, 2018.

27. Petr Danecek, Adam Auton, Goncalo Abecasis, Cornelis A. Albers, Eric Banks, Mark A. DePristo, Robert E. Handsaker, Gerton Lunter, Gabor T. Marth, Stephen T. Sherry, Gilean McVean, and Richard Durbin. The variant call format and vcftools. Bioinformatics, 27(15): 2156–2158, aug 2011. ISSN 1367-4803. doi: 10.1093/bioinformatics/btr330.

28. Petr Danecek, James K Bonfield, Jennifer Liddle, John Marshall, Valeriu Ohan, Martin O Pollard, Andrew Whitwham, Thomas Keane, Shane A McCarthy, Robert M Davies, and Heng Li. Twelve years of samtools and bcftools. GigaScience, 10(2), 02 2021. ISSN 2047-217X. doi: 10.1093/gigascience/giab008.giab008.

29. David H Alexander, John Novembre, and Kenneth Lange. Fast model-based estimation of ancestry in unrelated individuals. Genome Research, 19(9):1655–1664, 2009.

30. Thibaut Jombart, Sébastien Devillard, and François Balloux. Discriminant analysis of principal components: a new method for the analysis of genetically structured populations. BMC Genetics, 11(1):1–15, 2010.

31. Shaun Purcell, Benjamin Neale, Kathe Todd-Brown, Lori Thomas, Manuel AR Ferreira, David Bender, Julian Maller, Pamela Sklar, Paul IW De Bakker, Mark J Daly, and Pak C Sham. Plink: a tool set for whole-genome association and population-based linkage analyses. The American Journal of Human Genetics, 81(3):559–575, 2007.

32. Thibaut Jombart and Ismaïl Ahmed. adegenet 1.3-1: new tools for the analysis of genome-wide snp data. Bioinformatics, 27(21):3070–3071, 2011.

33. Brian J Knaus and Niklaus J Grünwald. vcfr: a package to manipulate and visualize variant call format data in r. Molecular Ecology Resources, 17(1):44–53, 2017.

34. Jonathan Terhorst, John A Kamm, and Yun S Song. Robust and scalable inference of population history from hundreds of unphased whole genomes. Nature Genetics, 49(2): 303–309, 2017.

35. Emanuel Schmid-Siegert, Namrata Sarkar, Christian Iseli, Sandra Calderon, Caroline Gouhier-Darimont, Jacqueline Chrast, Pietro Cattaneo, Frédéric Schütz, Laurent Farinelli, Marco Pagni, Michel Schneider, Jérémie Voumard, Michel Jaboyedoff, Christian Fankhauser, Christian S Hardtke, Laurent Keller, John R Pannell, Alexandre Reymond, Marc Robinson-Rechavi, Ioannis Xenarios, and Philippe Reymond. Low number of fixed somatic mutations in a long-lived oak tree. Nature Plants, 3(12):926–929, 2017.

36. R Core Team. R: A language and environment for statistical computing. 2021.

37. Desislava Petkova, John Novembre, and Matthew Stephens. Visualizing spatial population structure with estimated effective migration surfaces. Nature Genetics, 48(1):94–100, 2016.

38. Thorfinn Sand Korneliussen, Anders Albrechtsen, and Rasmus Nielsen. Angsd: analysis of next generation sequencing data. BMC Bioinformatics, 15(1):356, 2014.

39. Rasmus Nielsen, Thorfinn Korneliussen, Anders Albrechtsen, Yingrui Li, and Jun Wang. Snp calling, genotype calling, and sample allele frequency estimation from new-generation sequencing data. PloS One, 7(7):e37558, 2012.

40. Zhe Hou, Ang Li, and Jianguo Zhang. Genetic architecture, demographic history, and genomic differentiation of Populus davidiana revealed by whole-genome resequencing. Evolutionary Applications, 13(10):2582–2596, 2020.

41. Jarkko Salojärvi, Olli-Pekka Smolander, Kaisa Nieminen, Sitaram Rajaraman, Omid Safronov, Pezhman Safdari, Airi Lamminmäki, Juha Immanen, Tianying Lan, Jaakko Tanskanen, et al. Genome sequencing and population genomic analyses provide insights into the adaptive landscape of silver birch. Nature Genetics, 49(6):904–912, 2017.

42. Nikolaos Alachiotis and Pavlos Pavlidis. Raisd detects positive selection based on multiple signatures of a selective sweep and snp vectors. Communications Biology, 1(1):1–11, 2018.

43. Hadley Wickham. ggplot2: Elegant Graphics for Data Analysis. Springer-Verlag New York, 2016. ISBN 978-3-319-24277-4.

44. David M. Goodstein, Shengqiang Shu, Russell Howson, Rochak Neupane, Richard D. Hayes, Joni Fazo, Therese Mitros, William Dirks, Uffe Hellsten, Nicholas Putnam, and Daniel S. Rokhsar. Phytozome: a comparative platform for green plant genomics. Nucleic Acids Research, 40(D1):D1178–D1186, 2012.

45. Damian Smedley, Syed Haider, Steffen Durinck, Luca Pandini, Paolo Provero, James Allen, Olivier Arnaiz, Mohammad Hamza Awedh, Richard Baldock, Giulia Barbiera, Philippe Bardou, Tim Beck, Andrew Blake, Merideth Bonierbale, Anthony J Brookes, Gabriele Bucci, Iwan Buetti, Sarah Burge, Cédric Cabau, Joseph W. Carlson, Claude Chelala, Charalambos Chrysostomou, Davide Cittaro, Olivier Collin, Raul Cordova, Rosalind J. Cutts, Erik Dassi, Alex Di Genova, Anis Djari, Anthony Esposito, Heather Estrella, Eduardo Eyras, Julio Fernandez-Banet, Simon Forbes, Robert C Free, Takatomo Fujisawa, Emanuela Gadaleta, Jose M Garcia-Manteiga, David Goodstein, Kristian Gray, José Afonso Guerra-Assunção, Bernard Haggarty, Dong-Jin Han, Byung Woo Han, Todd Harris, Jayson Harshbarger, Robert K Hastings, Richard D Hayes, Claire Hoede, Shen Hu, Zhi-Liang Hu, Lucie Hutchins, Zhengyan Kan, Hideya Kawaji, Aminah Keliet, Arnaud Kerhornou, Sunghoon Kim, Rhoda Kinsella, Christophe Klopp, Lei Kong, Daniel Lawson, Dejan Lazarevic, Ji-Hyun Lee, Thomas Letellier, Chuan-Yun Li, Pietro Lio, Chu-Jun Liu, Jie Luo, Alejandro Maass, Jerome Mariette, Thomas Maurel, Stefania Merella, Azza Mostafa Mohamed, Francois Moreews, Ibounyamine Nabihoudine, Nelson Ndegwa, Céline Noirot, Cristian Perez-Llamas, Michael Primig, Alessandro Quattrone, Hadi Quesneville, Davide Rambaldi, James Reecy, Michela Riba, Steven Rosanoff, Amna Ali Saddiq, Elisa Salas, Olivier Sallou, Rebecca Shepherd, Reinhard Simon, Linda Sperling, William Spooner, Daniel M Staines, Delphine Steinbach, Kevin Stone, Elia Stupka, Jon W Teague, Abu Z Dayem Ullah, Jun Wang, Doreen Ware, Marie Wong-Erasmus, Ken Youens-Clark, Amonida Zadissa, Shi-Jian Zhang, and Arek Kasprzyk. The biomart community portal: an innovative alternative to large, centralized data repositories. Nucleic Acids Research, 43(W1):W589–W598, 2015.

46. Philippe Lamesch, Tanya Z Berardini, Donghui Li, David Swarbreck, Christopher Wilks, Rajkumar Sasidharan, Robert Muller, Kate Dreher, Debbie L Alexander, Margarita Garcia-Hernandez, Athikkattuvalasu S. Karthikeyan, Cynthia H Lee, William D Nelson, Larry Ploetz, Shanker Singh, April Wensel, and Eva Huala. The arabidopsis information resource (tair): improved gene annotation and new tools. Nucleic Acids Research, 40(D1):D1202–D1210, 2012.

47. Tanya Z Berardini, Suparna Mundodi, Leonore Reiser, Eva Huala, Margarita Garcia-Hernandez, Peifen Zhang, Lukas A Mueller, Jungwoon Yoon, Aisling Doyle, Gabriel Lander, Nick Moseyko, Danny Yoo, Iris Xu, Brandon Zoeckler, Mary Montoya, Neil Miller, Dan Weems, and Seung Y Rhee. Functional annotation of the arabidopsis genome using controlled vocabularies. Plant Physiology, 135(2):745–755, 2004.

48. Huaiyu Mi, Anushya Muruganujan, Xiaosong Huang, Dustin Ebert, Caitlin Mills, Xinyu Guo, and Paul D Thomas. Protocol update for large-scale genome and gene function analysis with the panther classification system (v. 14.0). Nature Protocols, 14(3):703–721, 2019.

49. David Q Bowen. Quaternary geology: a stratigraphic framework for multidisciplinary work. Pergamon Press Inc., 1978.

50. DF Belknap. Quaternary, April 2020.

51. Donald Edward Davis. The American chestnut: An environmental history. University of Georgia Press, 2021.

52. Kendra M Gurney, Paul G Schaberg, Gary J Hawley, and John B Shane. Inadequate cold tolerance as a possible limitation to american chestnut restoration in the northeastern united states. Restoration Ecology, 19(1):55–63, 2011.

53. Thomas M Saielli, Paul G Schaberg, Gary J Hawley, Joshua M Halman, and Kendra M Gurney. Nut cold hardiness as a factor influencing the restoration of american chestnut in northern latitudes and high elevations. Canadian Journal of Forest Research, 42(5):849–857, 2012.

54. Thomas M Saielli, Paul G Schaberg, Gary J Hawley, Joshua M Halman, and Kendra M Gurney. Genetics and silvicultural treatments influence the growth and shoot winter injury of american chestnut in vermont. Forest Science, 60(6):1068–1076, 2014.

55. Paul F Gugger, Sorel T Fitz-Gibbon, Ana Albarrán-Lara, Jessica W Wright, and Victoria L Sork. Landscape genomics of Quercus lobata reveals genes involved in local climate adaptation at multiple spatial scales. Molecular Ecology, 30(2):406–423, 2021.

56. Florian J Alberto, Sally N Aitken, Ricardo Alía, Santiago C González-Martínez, Heikki Hänninen, Antoine Kremer, François Lefèvre, Thomas Lenormand, Sam Yeaman, Ross Whetten, and Outi Savolainen. Potential for evolutionary responses to climate change–evidence from tree populations. Global Change Biology, 19(6):1645–1661, 2013.

57. Andrew J Eckert, Andrew D Bower, Santiago C González-Martínez, Jill L Wegrzyn, Graham Coop, and David B Neale. Back to nature: ecological genomics of loblolly pine (Pinus taeda, pinaceae). Molecular Ecology, 19(17):3789–3805, 2010.

58. Outi Savolainen and Tanja Pyhäjärvi. Genomic diversity in forest trees. Current Opinion in Plant Biology, 10(2):162–167, 2007.

59. JA Holliday, M Yuen, K Ritland, and SN Aitken. Postglacial history of a widespread conifer produces inverse clines in selective neutrality tests. Molecular Ecology, 19(18):3857–3864, 2010.

60. Jing Wang, Nathaniel R Street, Douglas G Scofield, and Pär K Ingvarsson. Natural selection and recombination rate variation shape nucleotide polymorphism across the genomes of three related Populus species. Genetics, 202(3):1185–1200, 2016.

61. Mary V Ashley, Saji T Abraham, Janet R Backs, and Walter D Koenig. Landscape genetics and population structure in valley oak (Quercus lobata née). American Journal of Botany, 102(12):2124–2131, 2015.

62. Samuel Royer-Tardif, Laura Boisvert-Marsh, Julie Godbout, Nathalie Isabel, and Isabelle Aubin. Appendices: Finding common ground: Toward comparable indicators of adaptive capacity of tree species to a changing climate. Ecology and Evolution, 11(19):13081–13100, 2021.

63. Gerald H Haug, Andrey Ganopolski, Daniel M Sigman, Antoni Rosell-Mele, George EA Swann, Ralf Tiedemann, Samuel L Jaccard, Jörg Bollmann, Mark A Maslin, Melanie J Leng, and Geoffrey Eglinton. North pacific seasonality and the glaciation of north america 2.7 million years ago. Nature, 433(7028):821–825, 2005.

64. Margaret B Davis. Quaternary history of deciduous forests of eastern north america and europe. Annals of the Missouri Botanical Garden, 70(3).

65. Paul A Delcourt. Goshen springs: late quaternary vegetation record for southern alabama. Ecology, 61(2):371–386, 1980.

66. William A Watts, Barbara CS Hansen, and Eric C Grimm. Camel lake: A 40 000-yr record of vegetational and forest history from northwest florida. Ecology, 73(3):1056–1066, 1992.

67. Paul A Delcourt, Hazel R Delcourt, Ronald C Brister, and Laurence E Lackey. Quaternary vegetation history of the mississippi embayment 1, 2. Quaternary Research, 13(1):111–132, 1980.

68. Hazel R Delcourt. Late quaternary vegetation history of the eastern highland rim and adjacent cumberland plateau of tennessee. Ecological Monographs, 49(3):255–280, 1979.

69. Jason S McLachlan, James S Clark, and Paul S Manos. Molecular indicators of tree migration capacity under rapid climate change. Ecology, 86(8):2088–2098, 2005.

70. Bowen S Crandall, GF Gravatt, and Margaret Milburn Ryan. Root disease of castanea species and some coniferous and broadleaf nursery stocks, caused by Phytophthora cinnamomi. Phytopathology, 35:162–80, 1945.

71. Jared W Westbrook, Joseph B James, Paul H Sisco, John Frampton, Sunny Lucas, and Steven N Jeffers. Resistance to Phytophthora cinnamomi in american chestnut (Castanea dentata) backcross populations that descended from two chinese chestnut (Castanea mollissima) sources of resistance. Plant Disease, 103(7):1631–1641, 2019.

72. Tetyana N Zhebentyayeva, Paul H Sisco, Laura L Georgi, Steven N Jeffers, M Taylor Perkins, Joseph B James, Frederick V Hebard, Christopher Saski, C Dana Nelson, and Albert G Abbott. Dissecting resistance to Phytophthora cinnamomi in interspecific hybrid chestnut crosses using sequence-based genotyping and qtl mapping. Phytopathology, 109(9):1594–1604, 2019.

73. M Taylor Perkins, Anna Claire Robinson, Martin L Cipollini, and J Hill Craddock. Identifying host resistance to Phytophthora cinnamomi in hybrid progeny of Castanea dentata and Castanea mollissima. HortScience, 54(2):221–225, 2019.

74. Abdelali Barakat, Denis S DiLoreto, Yi Zhang, Chris Smith, Kathleen Baier, William A Powell, Nicholas Wheeler, Ron Sederoff, and John E Carlson. Comparison of the transcriptomes of american chestnut (Castanea dentata) and chinese chestnut (Castanea mollissima) in response to the chestnut blight infection. BMC Plant Biology, 9(1):51, 2009.

75. Jared W Westbrook, Qian Zhang, Mihir K Mandal, Eric V Jenkins, Laura E Barth, Jerry W Jenkins, Jane Grimwood, Jeremy Schmutz, and Jason A Holliday. Optimizing genomic selection for blight resistance in american chestnut backcross populations: A trade-off with american chestnut ancestry implies resistance is polygenic. Evolutionary Applications, 13 (1):31–47, 2020.

76. Owen W Baughman, Alison C Agneray, Matthew L Forister, Francis F Kilkenny, Erin K Espeland, Rob Fiegener, Matthew E Horning, Richard C Johnson, Thomas N Kaye, Jeff Ott, and John B St. Clair. Strong patterns of intraspecific variation and local adaptation in great basin plants revealed through a review of 75 years of experiments. Ecology and Evolution, 9(11):6259–6275, 2019.

