## Supplemental Figures and Tables for "Whole-genome resequencing reveals the population structure, genomic diversity, and demographic history of American chestnut (*Castanea dentata*)"

### Supplementary

**immediate**

DRAFT

#### Supplement

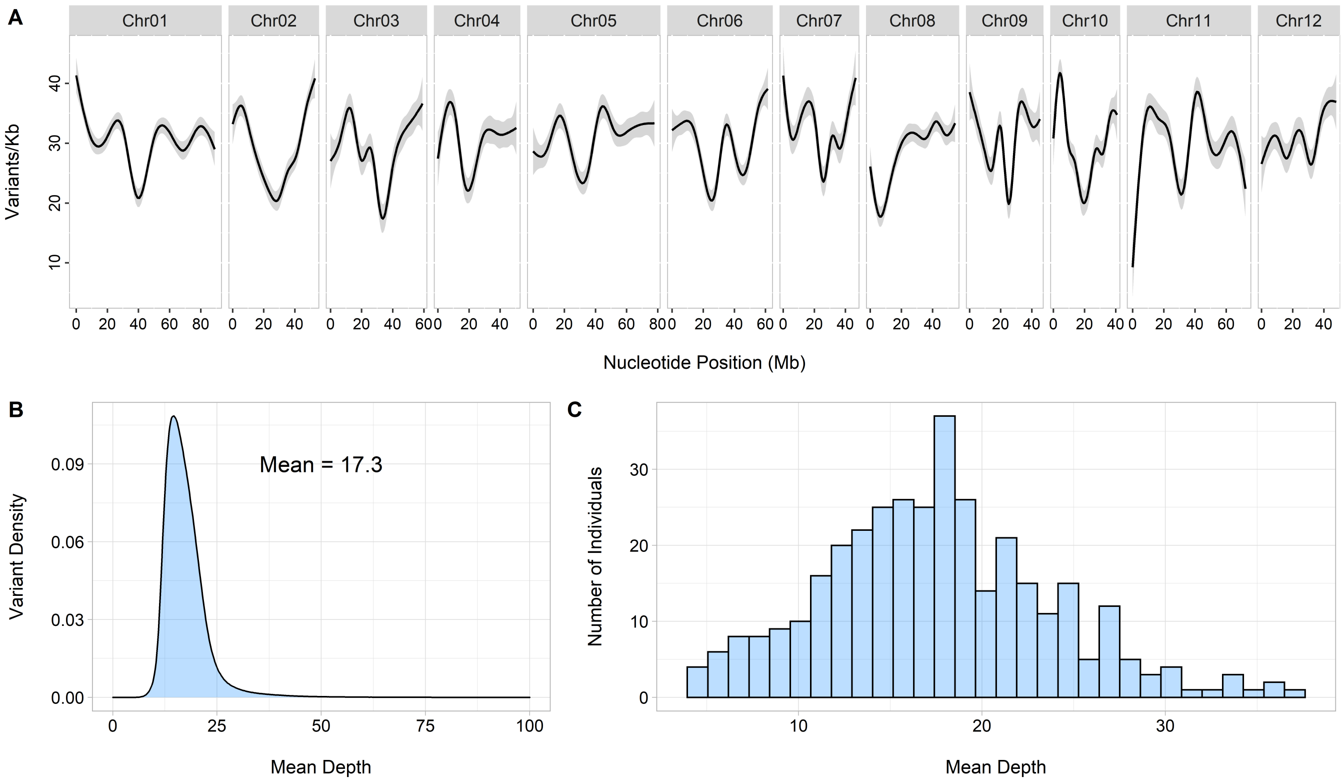


**Fig. S1.** Visualization of sequencing density and coverage in the filtered *Castanea dentata* genomic dataset. **A**. Genome-wide sequencing density. Greyed regions represent 95% confidence intervals. **B**. Mean depth per variant averaged across all samples. For visualization, 0.05% of all SNPs were excluded that had a coverage >100x. **C**. Mean depth per individual averaged across all variants.

DRAFT

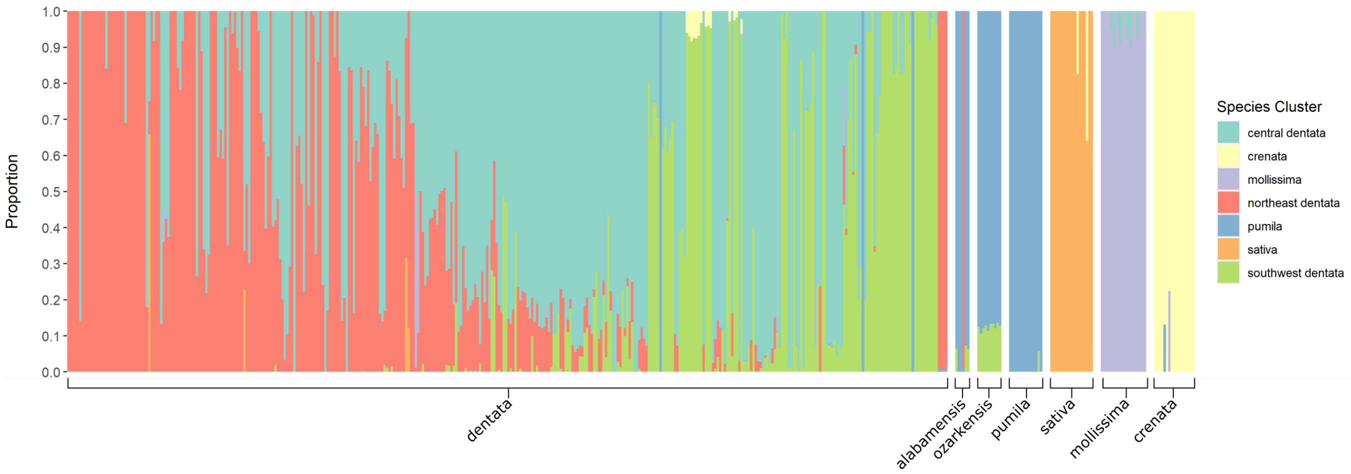


**Fig. S2.** ADMIXTURE analysis suggests that widespread hybridization does not occur within wild American chestnut populations. ADMIXTURE was performed with 384 American chestnut and 92 *Castanea* species samples. Each color bar represents the proportion of inferred species ancestry for a sample. K = 7 was determined to be the most appropriate cluster value.

DRAFT

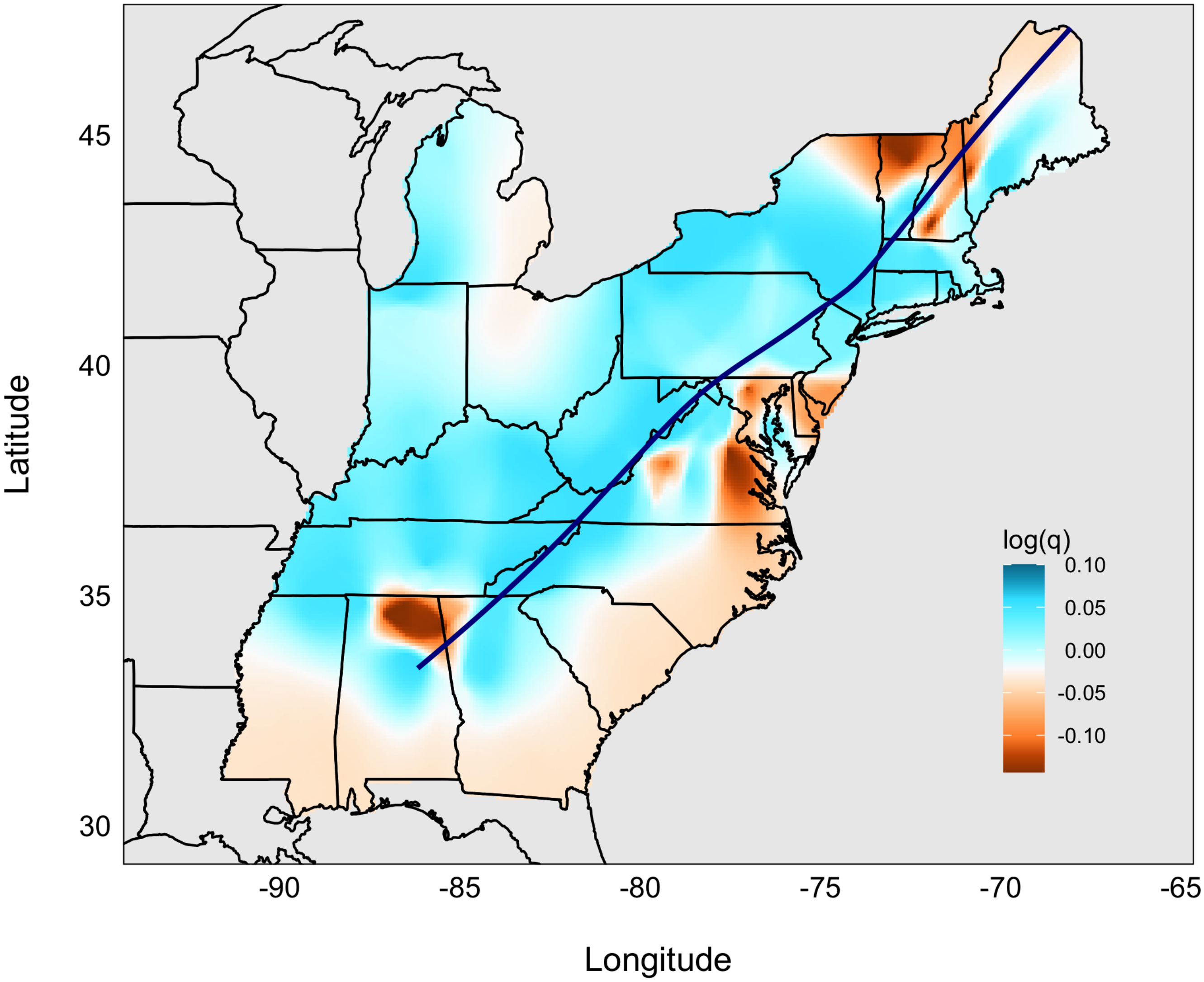


**Fig. S3.** Estimated genetic diversity within American chestnut throughout the eastern United States. The orange regions represent areas of lower than expected diversity, and the blue regions represent areas of higher than expected diversity. A loess line of Appalachian Mountain peaks was applied in *ggplot2* using peak locations obtained from https://en.wikipedia.org/wiki/List_of_mountains_of_the_Appalachians.

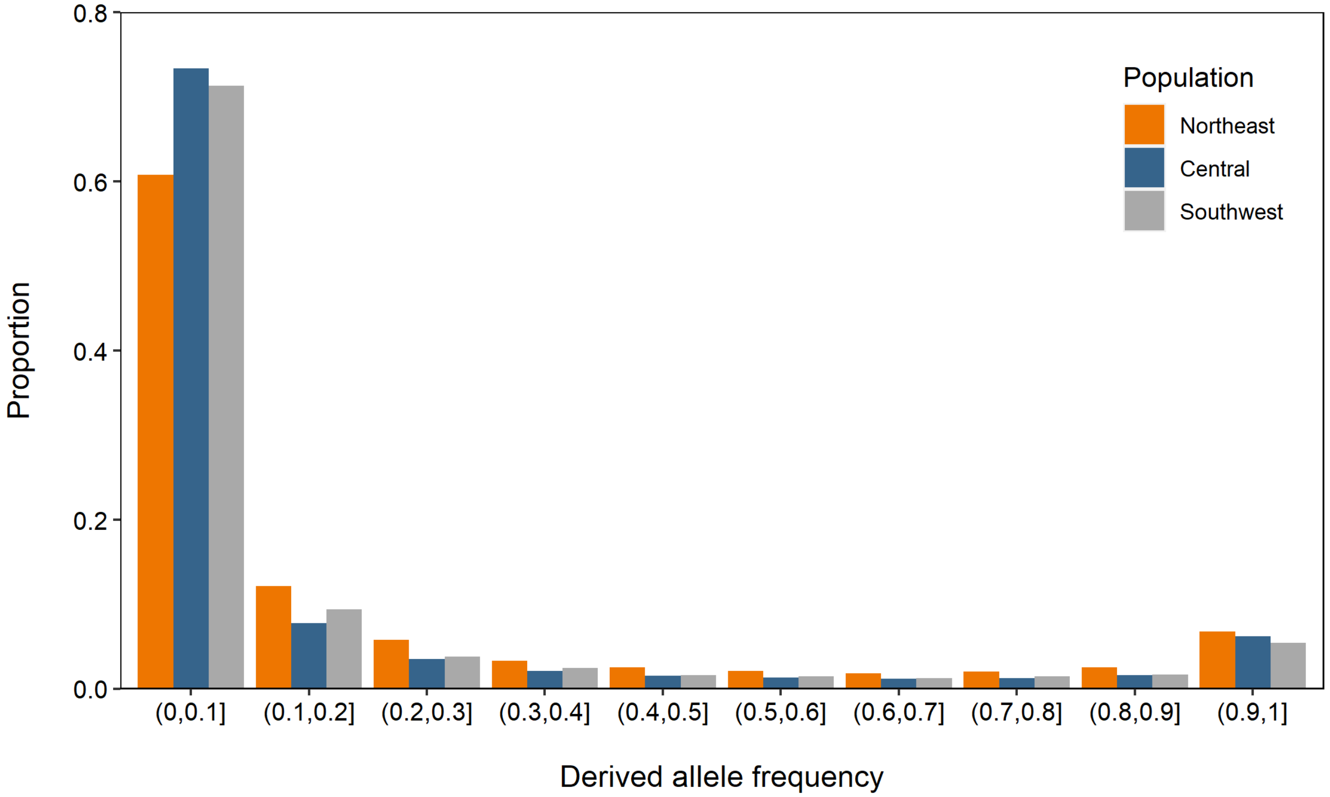


**Fig. S4.** Site-frequency-spectrum for each of the three wild American chestnut populations.

DRAFT

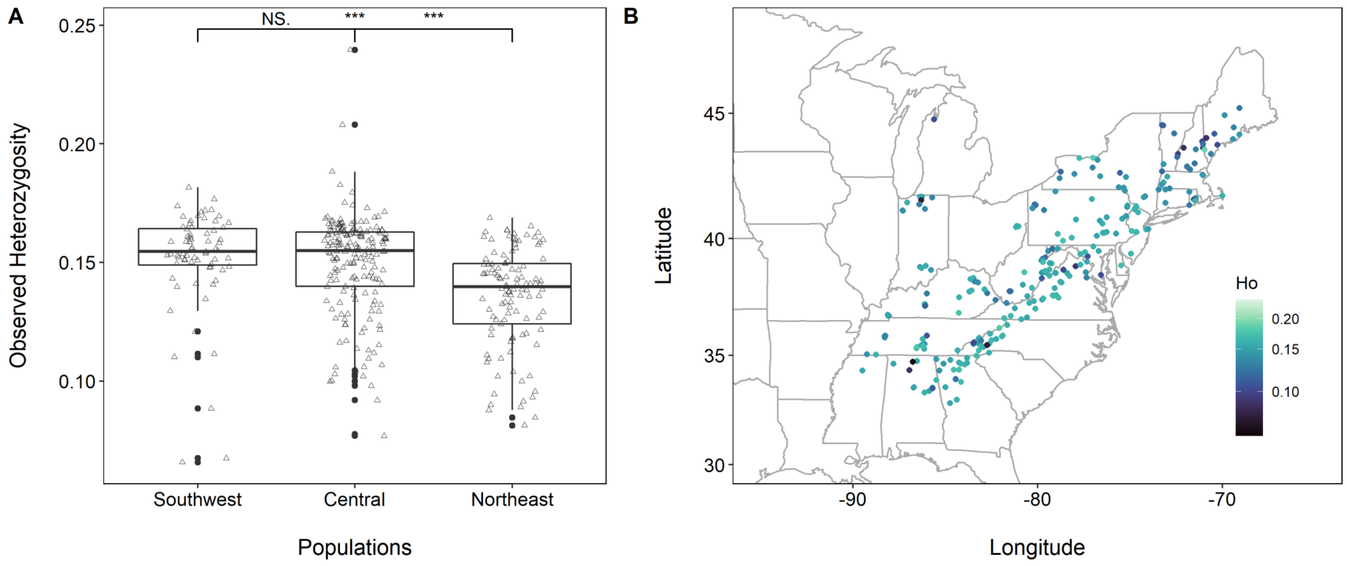


**Fig. S5.** Observed heterozygosity decreases as latitude increases between *Castanea dentata* populations. **A**. The northeast population has significantly less observed heterozygosity than the central and southwest populations ($p<0.001$, $p<0.001$). **B**. Map of sample locations and observed heterozygosity for each sample. Each circle represents a tree sample location and the color is the observed heterozygosity ratio

DRAFT

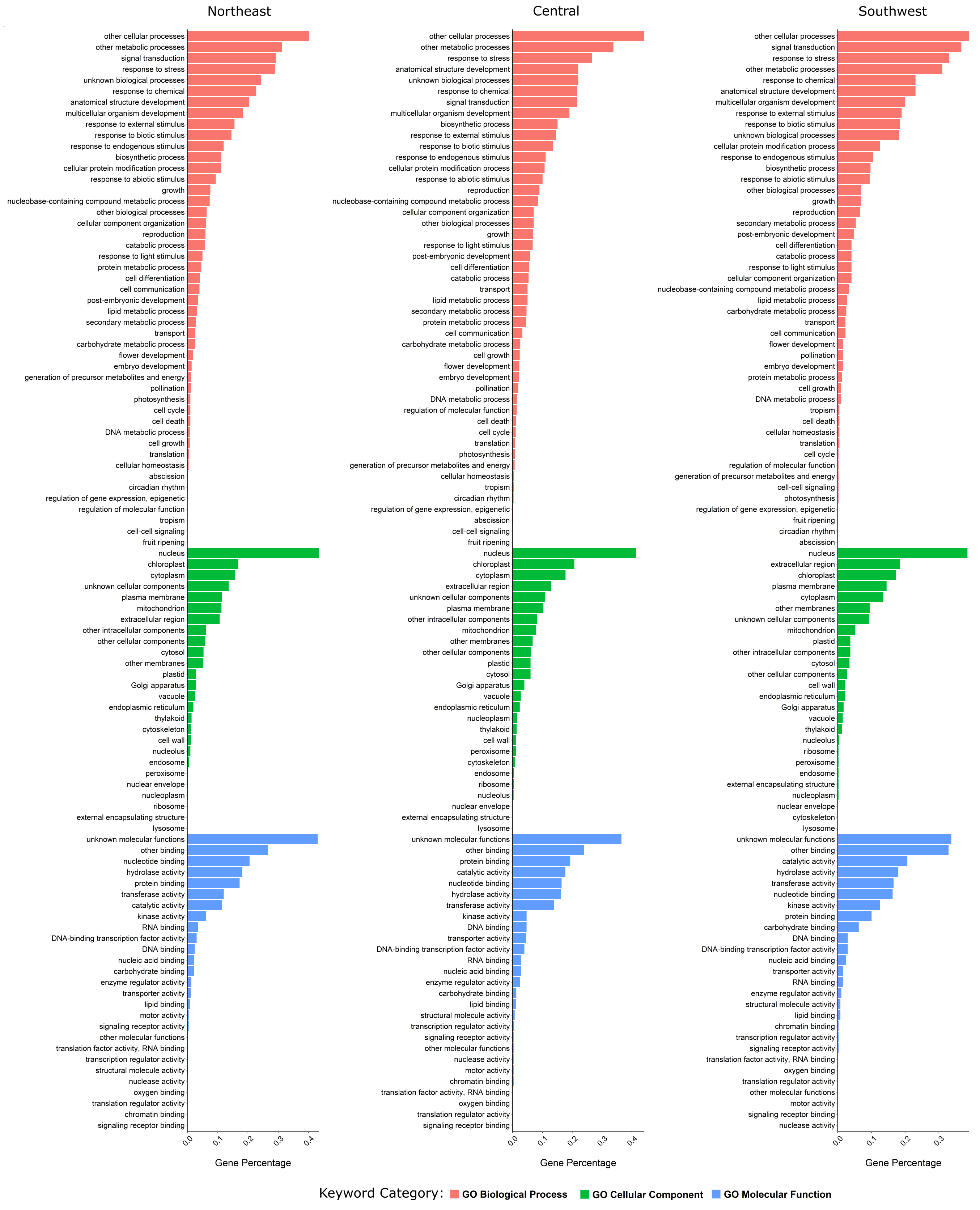


**Fig. S6.** GO annotations for all RAiSD outlier genes for each of the three American chestnut populations. The gene percentage was calculated by dividing the gene count for each category by its corresponding N value.

**Table S1.** GO enrichment analysis of the unique RAiSD outlier gene set for the southwest American chestnut population. The GO enrichment terms were identified as being significantly overrepresented for the GO biological process ($p<0.05$).

| GO ID | GO Description | Gene Count | Expected Count | Fold Enrichment | P-value |
|---|---|---|---|---|---|
| GO:0007166 | cell surface receptor signaling pathway | 20 | 0.38 | 53.28 | 1.64E-23 |
| GO:0050832 | defense response to fungus | 10 | 1.6 | 6.25 | 1.96E-02 |
| GO:0006468 | protein phosphorylation | 37 | 6.32 | 5.85 | 1.52E-14 |
| GO:0009620 | response to fungus | 12 | 2.19 | 5.48 | 8.69E-03 |
| GO:0016310 | phosphorylation | 38 | 6.94 | 5.47 | 4.31E-14 |
| GO:0007165 | signal transduction | 46 | 8.58 | 5.36 | 1.70E-17 |
| GO:0023052 | signaling | 47 | 8.82 | 5.33 | 7.32E-18 |
| GO:0007154 | cell communication | 51 | 10.39 | 4.91 | 3.12E-18 |
| GO:0006955 | immune response | 18 | 4.93 | 3.65 | 8.69E-03 |
| GO:0002376 | immune system process | 18 | 5 | 3.60 | 1.06E-02 |
| GO:0006796 | phosphate-containing compound metabolic process | 38 | 10.83 | 3.51 | 3.49E-08 |
| GO:0098542 | defense response to other organism | 17 | 4.89 | 3.47 | 3.12E-02 |
| GO:0006793 | phosphorus metabolic process | 38 | 11.11 | 3.42 | 7.22E-08 |
| GO:0051716 | cellular response to stimulus | 53 | 15.87 | 3.34 | 5.62E-12 |
| GO:0043207 | response to external biotic stimulus | 21 | 6.78 | 3.10 | 1.57E-02 |
| GO:0051707 | response to other organism | 21 | 6.78 | 3.10 | 1.57E-02 |
| GO:0009607 | response to biotic stimulus | 21 | 6.79 | 3.09 | 1.61E-02 |
| GO:0044419 | biological process involved in interspecies interaction between organisms | 21 | 6.86 | 3.06 | 1.85E-02 |
| GO:0036211 | protein modification process | 42 | 15.32 | 2.74 | 4.43E-06 |
| GO:0006464 | cellular protein modification process | 42 | 15.32 | 2.74 | 4.43E-06 |
| GO:0043412 | macromolecule modification | 42 | 17.32 | 2.42 | 1.76E-04 |
| GO:0044267 | cellular protein metabolic process | 43 | 20.83 | 2.06 | 9.14E-03 |
| GO:0050896 | response to stimulus | 69 | 33.69 | 2.05 | 2.24E-06 |
| GO:0019538 | protein metabolic process | 44 | 21.77 | 2.02 | 1.33E-02 |
| GO:0050794 | regulation of cellular process | 55 | 29.16 | 1.89 | 5.08E-03 |

DRAFT

**Table S2.** GO enrichment analysis of the unique RAiSD outlier gene set for the northeast American chestnut population. The GO enrichment terms were identified as being significantly overrepresented for the GO biological process ($p<0.05$).

| GO ID | GO Description | Gene Count | Expected Count | Fold Enrichment | P-value |
|---|---|---|---|---|---|
| GO:1900459 | positive regulation of brassinosteroid mediated signaling pathway | 4 | 0.06 | 69 | 3.51E-03 |
| GO:0007166 | cell surface receptor signaling pathway | 8 | 0.44 | 18.1 | 1.02E-04 |
| GO:0002376 | immune system process | 24 | 5.89 | 4.07 | 2.72E-05 |
| GO:0006955 | immune response | 23 | 5.8 | 3.96 | 9.27E-05 |
| GO:0098542 | defense response to other organism | 22 | 5.76 | 3.82 | 3.51E-04 |
| GO:0007165 | signal transduction | 35 | 10.1 | 3.46 | 5.36E-07 |
| GO:0023052 | signaling | 35 | 10.38 | 3.37 | 1.10E-06 |
| GO:0007154 | cell communication | 41 | 12.23 | 3.35 | 3.02E-08 |
| GO:0006952 | defense response | 22 | 7.28 | 3.02 | 1.53E-02 |
| GO:0051716 | cellular response to stimulus | 46 | 18.69 | 2.46 | 3.45E-05 |

DRAFT

**Table S3.** GO enrichment analysis of the unique RAiSD outlier gene set for the central American chestnut population. The GO enrichment terms were identified as being significantly overrepresented for the GO biological process ($p<0.05$).

| GO ID | GO Description | Gene Count | Expected Count | Fold Enrichment | P-value |
|---|---|---|---|---|---|
| GO:0050896 | response to stimulus | 84 | 54.02 | 1.56 | 4.89E-02 |

DRAFT

**Table S4.** GO enrichment analysis of the shared RAiSD outlier gene set for three American chestnut populations. The GO enrichment terms were identified as being significantly underrepresented or overrepresented for the GO biological process ($p<0.05$).

| GO ID | GO Description | Gene Count | Expected Count | Fold Enrichment | P-value |
|---|---|---|---|---|---|
| GO:0050832 | defense response to fungus | 5 | 0.3 | 16.5 | 3.73E-02 |
| GO:0008150 | biological_process | 13 | 24.72 | 0.53 | 2.64E-02 |

DRAFT
